# Optimizing predictive models to prioritize viral discovery in zoonotic reservoirs

**DOI:** 10.1101/2020.05.22.111344

**Authors:** Daniel J. Becker, Gregory F. Albery, Anna R. Sjodin, Timothée Poisot, Laura M. Bergner, Tad A. Dallas, Evan A. Eskew, Maxwell J. Farrell, Sarah Guth, Barbara A. Han, Nancy B. Simmons, Michiel Stock, Emma C. Teeling, Colin J. Carlson

## Abstract

Despite global investment in One Health disease surveillance, it remains difficult—and often very costly—to identify and monitor the wildlife reservoirs of novel zoonotic viruses. Statistical models can be used to guide sampling prioritization, but predictions from any given model may be highly uncertain; moreover, systematic model validation is rare, and the drivers of model performance are consequently under-documented. Here, we use bat hosts of betacoronaviruses as a case study for the data-driven process of comparing and validating predictive models of likely reservoir hosts. In the first quarter of 2020, we generated an ensemble of eight statistical models that predict host-virus associations and developed priority sampling recommendations for potential bat reservoirs and potential bridge hosts for SARS-CoV-2. Over more than a year, we tracked the discovery of 40 new bat hosts of betacoronaviruses, validated initial predictions, and dynamically updated our analytic pipeline. We find that ecological trait-based models perform extremely well at predicting these novel hosts, whereas network methods consistently perform roughly as well or worse than expected at random. These findings illustrate the importance of ensembling as a buffer against variation in model quality and highlight the value of including host ecology in predictive models. Our revised models show improved performance and predict over 400 bat species globally that could be undetected hosts of betacoronaviruses. Although 20 species of horseshoe bats (*Rhinolophus* spp.) are known to be the primary reservoir of SARS-like viruses, we find at least three-fourths of plausible betacoronavirus reservoirs in this bat genus might still be undetected. Our study is the first to demonstrate through systematic validation that machine learning models can help optimize wildlife sampling for undiscovered viruses and illustrates how such approaches are best implemented through a dynamic process of prediction, data collection, validation, and updating.

## Introduction

Identifying likely reservoirs of zoonotic pathogens is challenging^1^. Sampling wildlife for the presence of active or recent infection (i.e., seropositivity) represents the first stage of a pipeline for proper inference of host species^2^, but sampling is often limited in phylogenetic, temporal, and spatial scale by logistical constraints^3^. Given such restrictions, statistical models can play a critical role in helping to prioritize pathogen surveillance by narrowing the set of plausible sampling targets, by either ruling out clades of low-likelihood hosts^4,5^ or predicting high-risk clades^6^. For example, machine learning approaches have generated candidate lists of likely, but unsampled, primate reservoirs of Zika virus, bat reservoirs of filoviruses, and avian reservoirs of *Borrelia burgdorferi*^*7–9*^.

At the same time, host predictions are rarely validated empirically^10^. Occasional case studies suggest both success and failures. For example, models predicted *Eonycteris spelaea* as an undetected bat host of filoviruses^7^, which was later confirmed through field sampling in southeast Asia^11,12^. Similarly, models of mosquito–Zika virus interactions predicted *Culex quinquefasciatus* as a likely vector^13^, which was rapidly validated through experimental competence trials^14,15^. A recent model of Nipah virus in India also predicted several bat species as undetected hosts^2^. However, experimental infection of the predicted *Rousettus aegyptiacus* demonstrated that this species cannot support virus replication^16^. Further, Nipah virus was recently found circulating in *Pipistrellus pipistrellus*, a species with a low predicted probability of being a host^17^. More generally, predictions from most models remain either untested or opportunistically validated, limiting insight into which approaches have greatest predictive accuracy. Systematically validating predictions would provide critical insights into the broader utility (or inefficacy) of different models in zoonosis research. Moreover, these modeling approaches are generally developed in isolation; implementation of multiple modeling approaches collaboratively and simultaneously, as part of a model-to-validation workflow, could reduce redundancy and apparent disagreement at the earliest stages of pathogen tracing while advancing predictive analytics by addressing inter-model reliability.

Coronaviruses (CoVs) are an ideal family of viruses with which to compare and validate predictive models of likely zoonotic reservoirs. CoVs are positive-sense, single-stranded RNA viruses that are detected across mammals and birds^18^. They have a broad host range, a high mutation rate, and the largest genomes of any RNA viruses, but they have also evolved mechanisms for RNA proofreading and repair to mitigate the deleterious effects of a high recombination rate acting over a large genome^19^. Consequently, CoVs fit the profile of viruses with high zoonotic potential. There are eight human CoVs (three and five in the genera *Alphacoronavirus* and *Betacoronavirus*, respectively), of which three are highly pathogenic in humans: SARS-CoV, MERS-CoV, and SARS-CoV-2. These are zoonotic and widely agreed to have evolutionary origins in bats^20–23^.

Our collective experience with both SARS-CoV and MERS-CoV illustrates the difficulty of tracing specific animal hosts of emerging viruses. During the 2002–2003 SARS epidemic, SARS-CoV was traced to the masked palm civet (*Paguma larvata*)^24^, but the ultimate origin remained unknown for several years. Horseshoe bats (family Rhinolophidae: genus *Rhinolophus*) were implicated as reservoir hosts in 2005, but their SARS-like CoVs were not identical to circulating human strains^21^. Stronger evidence from 2017 placed the most likely evolutionary origin of SARS-CoV in *Rhinolophus ferrumequinum* or *R. sinicus*^*25*^. Presently, there is even less certainty about the origins of MERS-CoV, although spillover to humans occurs relatively often through contact with dromedary camels (*Camelus dromedarius*). A virus with 100% nucleotide identity in a ∼200 base pair region of the MERS-CoV polymerase gene was detected in *Taphozous perforatus* (family Emballonuridae) in Saudi Arabia^26^; however, based on spike gene similarity, other sources treat HKU4 virus from *Tylonycteris pachypus* (family Vespertilionidae) in China as the most closely related bat virus^27,28^. Several bat CoVs have shown close phylogenetic relationships to MERS-CoV, with a surprisingly broad geographic distribution from Mexico to China^29–32,^.

Coronavirus disease 2019 (COVID-19) is caused by SARS-CoV-2, a novel virus with presumed evolutionary origins in bats. Although the earliest cases were linked to a wildlife market^23^, contact tracing was limited, and there has been no definitive identification of the wildlife contact that resulted in spillover nor a true “index case.” The divergence time between SARS-CoV-2 and two of the closest-related bat viruses (RaTG13 from *Rhinolophus affinis* and RmYN02 from *Rhinolophus malayanus*) has been estimated as 40-50 years^33^, suggesting that the main host(s) involved in spillover remain unknown. Evidence of viral recombination in pangolins has been proposed but is unresolved^33^. SARS-like betacoronaviruses have been recently isolated from Sunda pangolins (*Manis javanica*) traded in wildlife markets^34,35^, and these viruses have a very high amino acid identity to SARS-CoV-2, but only show a ∼90% nucleotide identity with SARS-CoV-2 or Bat-CoV RaTG13^36^. None of these host species are universally accepted as the origin of SARS-CoV-2 nor are any of the viruses a clear SARS-CoV-2 progenitor, and a “better fit” wildlife reservoir could likely still be identified. However, substantial gaps in betacoronavirus sampling across wildlife limit actionable inference about plausible reservoir hosts and bridge hosts for SARS-CoV-2^37^.

### Building a predictive ensemble

Here, we use betacoronaviruses in bats as a case study for the data-driven process of comparing and validating predictive models of likely reservoir hosts, with the ultimate aim to help prioritize surveillance for known and future zoonotic viruses. We focused on betacoronaviruses rather than SARS-like CoVs (subgenus *Sarbecovirus*) specifically, as the latter are only characterized from a very small number of bat species in publicly available data. This sparsity makes current modeling methods poorly suited to more precisely infer potential reservoir hosts of sarbecoviruses specifically. Instead, we used predictive models to (1) identify bats (and other mammals) that may broadly host any betacoronavirus and (2) identify species with a high viral sharing probability with the two *Rhinolophus* species carrying the earliest known close viral relatives of SARS-CoV-2. In mid-2020, at the early stages of the COVID-19 pandemic, we developed a standardized dataset of mammal–virus associations by integrating a previously published edgelist^38^ with a targeted scrape of all GenBank accessions for Coronaviridae and their associated hosts. Our final dataset spanned 710 host species and 359 virus genera, including 107 mammal hosts of betacoronaviruses as well as hundreds of other (non-CoV) association records. We integrated our host– virus data with a mammal phylogenetic supertree^39^ and over 60 standardized ecological traits of bat species^7,40,41^.

We then used these data to generate an ensemble of predictive models and drew on two popular approaches to identify candidate bat reservoir hosts of betacoronaviruses (Table 1). Network-based methods estimate a full set of “true” unobserved host–virus interactions based on a recorded network of associations (here, pairs of host species and associated viral genera). These methods are increasingly popular as a way to identify latent processes structuring ecological networks^42–44^, but they are often confounded by sampling bias and often can only make predictions for species within the observed network (i.e., those that have available virus data; in-sample prediction). In contrast, trait-based methods use observed relationships concerning host traits to identify species that fit the morphological, ecological, or phylogenetic profile of known host species of a given pathogen, and rank the suitability of unknown hosts based on these trait characteristics^8,45^. These methods may be more likely to recapitulate patterns in observed host–pathogen association data (e.g., geographic biases in sampling, phylogenetic similarity in host morphology), but they more easily correct for sampling bias and can predict host species without known viral associations (i.e., out-of-sample prediction).

**Table 1.** Scope and calibration of different predictive modeling approaches. Some methods use pseudoabsences to expand the scale of prediction but still only analyze existing host–virus data, with no out-of-sample inference, whereas other methods can predict freshly onto new data.

| **Model approach** | **Prediction on hosts without known associations (out-of-sample)** | **Predictive extent and use of pseudoabsences** |
| --- | --- | --- |
| **Network-based 1**<br>k-Nearest neighbors | No | Only predicts link probabilities among species in the association data |
| **Network-based 2**<br>Linear filter | No | Only predicts link probabilities among species in the association data |
| **Network-based 3**<br>Plug and play | No | Uses pseudoabsences to predict over all mammals in association data, using latent approach |
| **Network-based 4**<br>Scaled-phylogeny | No | Only predicts link probabilities among species in the association data |
| **Trait-based 1**<br>Boosted regression trees | Yes | Uses pseudoabsences for all bats in trait data to predict over all species, including those without known associations |
| **Trait-based 2**<br>Bayesian additive regression trees | Yes | Uses pseudoabsences for all bats in trait data to predict over all species, including those without known associations |
| **Trait-based 3**<br>Neutral phylogeographic | Yes | Trains on a broader network, and predicts sharing probabilities among any mammals in phylogeny and IUCN range map data |
| **Hybrid 1**<br>Two-step kernel ridge regression | Yes | Uses pseudoabsences for all bats in trait data to predict over all species, including those without known associations |

In total, we implemented eight different predictive models of host-virus associations, including four network-based approaches, three trait-based approaches, and one hybrid approach using trait and phylogenetic information to make network predictions. These efforts generated eight ranked lists of suspected bat hosts of betacoronaviruses. Each ranked list was then scaled proportionally and consolidated in an ensemble of recommendations for betacoronavirus sampling and broader eco-evolutionary research. Next, approximately one year after our initial model ensemble, we reran our entire analytic pipeline with new bat betacoronavirus detections, taking advantage of the recent proliferation of published research on bat CoVs. This provided an unprecedented opportunity to rapidly compare model performance, provide up-to-date predictions of likely but unsampled bat hosts, and critically assess model accuracy in the context of ongoing sampling for bat CoVs.

### Predicted bat reservoirs of betacoronaviruses

Our initial ensemble found wide variation in model performance; individual models explained 0–69% of the variance in betacoronavirus positivity (mean of 25%), whereas the ensemble generally had improved predictive capacity (*R*^*2*^ = 42%; Supplemental Figure 1). The predictions of bat betacoronavirus hosts derived from network- and trait-based modeling approaches displayed strong inter-model agreement within each group but largely differed between groups (Figure 1). Of the 1,037 included bat species not known to be infected by betacoronaviruses during our initial analysis in 2020 (against 79 known positives), our models identified between 7 and 722 potential hosts based on a 10% omission threshold (90% sensitivity). Applying this same threshold to our ensemble predictions, our initial models identified 371 bat species as likely undetected hosts. Importantly, only 48 suspect hosts were identified in-sample, whereas we identified 323 suspect hosts out-of-sample, highlighting that most undiscovered hosts—and in turn undiscovered betacoronaviruses—should be in unsampled bat species.

**Figure 1.**
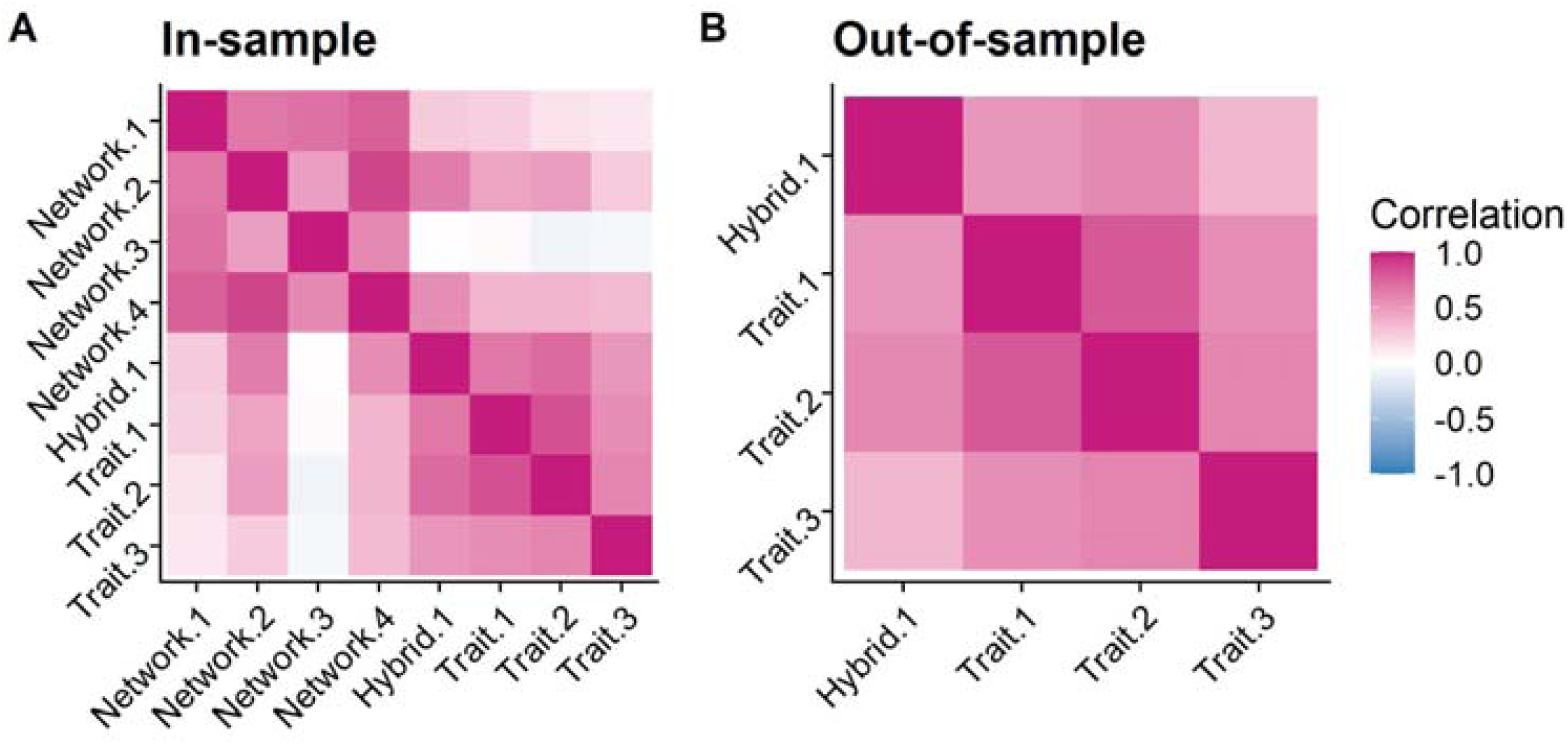
Broad agreement across an ensemble of predictive modeling approaches. The pairwise Spearman’s rank correlations between models’ ranked species-level predictions were generally substantial and positive. Models are arranged in decreasing order of their mean correlation with other models. Models that used trait data made more similar predictions to each other than to approaches using network methods with the same data (A); network-based models that used some ecological data made more similar predictions to all other models (Network-4, which uses phylogeny, and Hybrid-1, which uses both phylogeny and trait data). All models that could make out-of-sample predictions used trait data (B) and showed strong agreement.

This multi-model ensemble predicted undiscovered betacoronavirus bat hosts with striking geographic and taxonomic patterning (Figure 2). In-sample, predicted hosts were globally distributed and recapitulated geographic patterns of known bat betacoronavirus hosts in Europe, the Neotropics, and southeast Asia; however, our models also predicted high richness of likely bat reservoirs in North America. Applying a graph partitioning algorithm (phylogenetic factorization) to the bat phylogeny^46^, we similarly found that both betacoronavirus positivity and in-sample predictions were, on average, lowest for the superfamilies Noctilionoidea and Vespertilionoidea in the Yangochiroptera. This makes intuitive sense, as these taxa do not include the groups known to harbor the vast majority of betacoronaviruses detected in bats (e.g., *Rhinolophus*, Hipposideridae). In contrast, our out-of-sample predicted hosts were more starkly clustered in much of sub-Saharan Africa and southeast Asia (e.g., Vietnam, Myanmar, and southern China), with no representation in the western hemisphere. Likewise, out-of-sample predictions were lower in Neotropical bat families (e.g., Noctilionidae, Mormoopidae, Phyllostomidae), most emballonurids, and primarily Neotropical molossids, whereas the *Rhinolophus* genus and most of the Old World subfamily Pteropodinae were predicted to be more likely to host betacoronaviruses (Supplemental Table 1).

**Figure 2.**
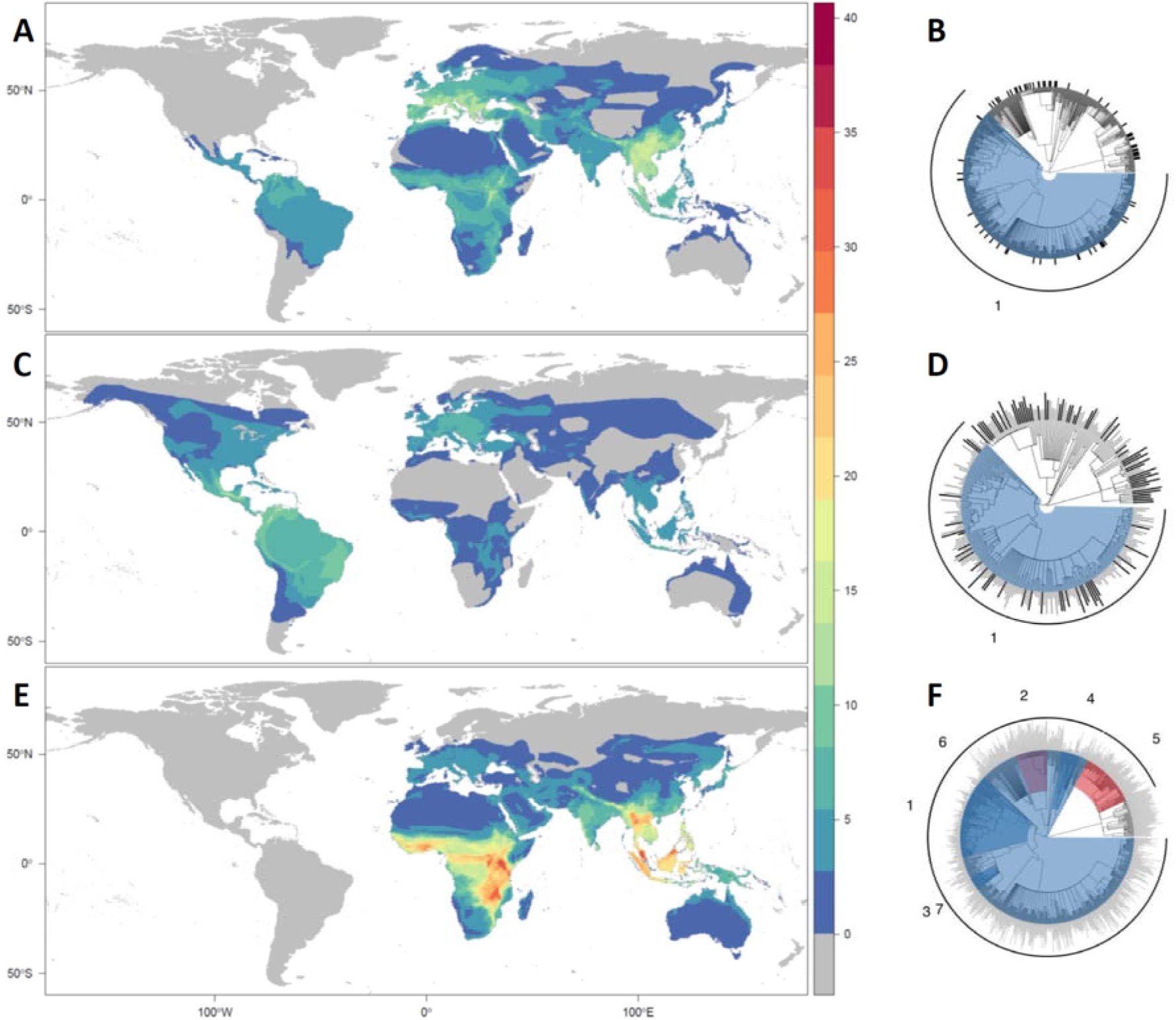
The geographic and evolutionary distribution of known and predicted bat hosts of betacoronaviruses (initial ensemble predictions). Known hosts of betacoronaviruses (A, B) are found worldwide, but particularly in southern Asia and southern Europe. Taxonomically, betacoronaviruses are less common in two superfamilies of the Yangochiroptera, the Noctilionoidea and Vespertilionoidea (clade 1). The predicted in-sample bat hosts (i.e., those with any viral association records; C, D) tend to recapitulate observed geographic patterns of known hosts but with a higher concentration in the Neotropics. Similarly, taxonomic patterns reflect those of known betacoronavirus hosts. In contrast, the out-of-sample bat host predictions based on phylogeny and ecological traits (E, F) are mostly clustered in Myanmar, Vietnam, and southern China, with none in the Neotropics or North America. Predicted hosts are likewise more common in the Rhinolophidae (clade 2) and subfamilies of Old World bats (clade 5) and are rare in many Neotropical taxa (clades 1 and 7) and emballanurids (clades 3 and 4). In the phylogenies, bar height indicates betacoronavirus positivity (B) or predicted rank (D, F; higher values indicate lower proportional ranks). Colors indicate clades identified through phylogenetic factorization (red indicates clades more likely to contain hosts, blue indicates less likely hosts; see Supplemental Table 1 for more information).

Because only trait-based models were capable of out-of-sample prediction, the differences in geographic and taxonomic patterns of our predictions likely reflect distinctions between the network- and trait-based modeling approaches. We suggest these should be considered as qualitatively different lines of evidence. Network approaches proportionally upweight species with high observed viral diversity, recapitulating sampling biases largely unrelated to coronaviruses (e.g., frequent screening for rabies lyssaviruses in the common vampire bat *Desmodus rotundus*, which has been sampled in a comparatively limited capacity for coronaviruses^31,47–49^). Highly ranked species may also have been previously sampled without evidence of betacoronavirus presence; for example, *Rhinolophus luctus* and *Macroglossus sobrinus* from China and Thailand, respectively, tested negative for betacoronaviruses, but detection probability was limited by small sample sizes^50–52^. In contrast, trait-based approaches are constrained by their reliance on phylogeny, ecological traits, and geographic covariates, all of which made models more likely to recapitulate existing spatial (i.e., clustering in southeast Asia) and taxonomic (i.e., the *Rhinolophus* genus) patterns. However, out-of-sample predictions are, by definition, inclusive of unsampled bat hosts^53^, which potentially offer greater return on viral discovery investment.

### Model validation

Following this initial 2020 model ensemble, we used broad literature searches to systematically track betacoronavirus-positive bat species that were missed in our initial data compilation (e.g., CoV sequences that were not annotated to genus on GenBank^52,54^). These searches also tracked the exponential increase in data on bat CoVs stemming from the emergence of SARS-CoV-2 that were published after our first model ensemble. A year after our initial data compilation (June 2021), we also reran our initial scrape of GenBank to identify new betacoronavirus-positive bats, limiting our search to matches to *Betacoronavirus* (taxid: 694002) and Chiroptera (taxid: 9397); however, this did not recover any additional positive host species not already recorded as positives in our updated data. We also mined publically available metagenomic and transcriptomic datasets for evidence of betacoronavirus infection^55–57^. However, no published libraries contained evidence of betacoronaviruses (see Supplement). Lastly, we analyzed the wildlife testing data from the USAID Emerging Pandemic Threats PREDICT program collected from 2009-2019, which was publicly released in June 2021 and includes a number of betacoronaviruses that were discovered during the program’s run (2009 to 2019) but have only published and identified down to the genus level in the full release.

In total, we uncovered 40 novel bat hosts of betacoronaviruses that were either absent from our original dataset or newly discovered after our initial analyses. This data update resulted in a total of 119 positive bat species, and we continue to collate these records (https://www.viralemergence.org/betacov). Of these 40 new hosts, the original ensemble correctly predicted only 24 (60% success rate), but some submodels performed significantly better than others, including both network- and trait-based methods: Network-1identified 20 of the 21 in-sample novel hosts (95%), while Trait-1 identified 37 of the 40 novel hosts (92.5%). The high performance of both of these top models, and their high performance on the training data (Supplemental Figure 1), suggest both approaches contributed usefully to the initial ensemble.

The 40 newly discovered hosts also enabled us to develop a new kind of performance metric for machine learning tasks with presence-only validation data (i.e., new “positives” can be collected, while “negatives” are substantially more difficult to prove). If a model makes predictions at random, the predicted prevalence of positives in the training data should be roughly the same as the success rate with novel test data. For example, a “coin toss” model will estimate that approximately 50% of species are betacoronavirus hosts and would likewise successfully identify approximately 50% of newly discovered hosts. A high-performing model, on the other hand, will identify a higher proportion of newly discovered hosts than expected at random. To evaluate how models perform in this regard, we developed a new diagnostic called the training prevalence-test sensitivity curve (TPTSC) that can be applied to modeling problems where training data are composed of a mix of true positives, true negatives, and false negatives, but test data only include novel true positives (Figure 3). The TPTSC plots the assumed prevalence in the training data against the sensitivity in the test data at every possible threshold from 0 to 1; these curves can be treated like receiver-operator or precision-recall curves, where a higher area under the curve (AUC) indicates better-than-random performance. Using the AUC-TPTSC scores, we found that trait-based and hybrid models consistently performed extremely well (Trait-1: 0.80; Trait-2: 0.79; Trait-3: 0.73; Hybrid-1: 0.68), while network methods performed at-random or worse (Network-1: 0.58; Network-2: 0.43; Network-3: 0.51; Network-4: 0.55). Accordingly, the ensemble performed comparably to the trait-based models (AUC-TPTSC = 0.75).

**Figure 3.**
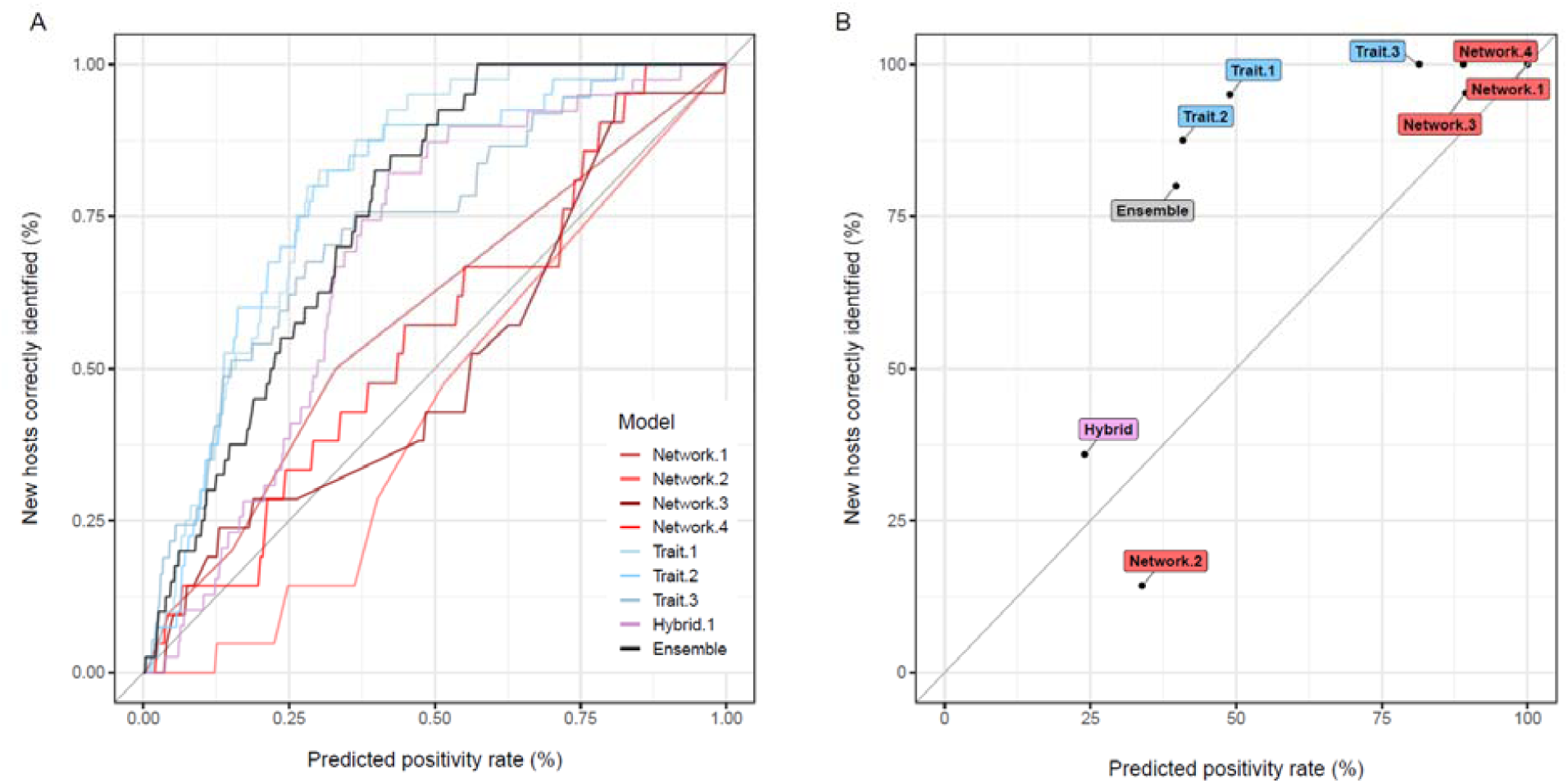
Measuring model performance with novel data. Performance is based on the comparison of total predicted prevalence (i.e., how many species are predicted positives?) with the sensitivity measured from validation data (i.e., how many of the 40 new species are correctly identified?). The null expectation for a model with random performance is these should be equivalent (e.g., a coin toss model will say that 50% of all bats have a betacoronavirus, and will be right 50% of the time), while a model with strong performance will be above that null expectation (*thin grey line*). (A) The training prevalence-test sensitivity curve (TPTSC) is a novel diagnostic that is conceptually similar to the receiver-operator curve (ROC), in that the model is evaluated at each possible scaled rank threshold between 0 and 1. (B) The same analysis, but only showing the point estimate of positivity created by each model’s internally-calibrated threshold. For model-guided sampling, the best model would be one that predicts a low-to-medium positivity rate and has a disproportionately high sensitivity (i.e., in the upper left corner). Both (A) and (B) show that the trait-based models (including the hybrid model) perform well, while the network-only models perform roughly at-random or worse than random (i.e., close to the line); the ensemble model, which includes all eight, performs comparably to the two best trait models and better than six of the eight component models.

These results have two key implications for future efforts to target sampling for putative reservoir hosts. First, ensembling can be useful as a buffer against variation in model quality, particularly in settings when the underlying drivers of model performance have yet to be identified. Second, and perhaps more importantly, models failed to have better-than-random performance without trait data that characterized bat ecology, even when they included phylogenetic data (e.g., Network-4). Part of this difference may also be attributable to the different scope of prediction: the response variable of trait-based models is betacoronavirus presence, while betacoronavirus-relevant predictions were extracted from a broader set of predictions made by the network models. However, this is contraindicated by the results of Hybrid-1, which performed comparably to the other trait-based models. Therefore, we conclude that making meaningful predictions about likely zoonotic reservoirs is best accomplished by incorporating detailed information on host ecology. The vastly greater performance of trait-based models provides another compelling reason, in addition to other One Health and conservation rationales, to better understand the fundamental ecology and evolution of bats.

### Dynamic prediction

Inclusion of these 40 novel bat hosts dramatically improved the performance of our predictive models. When revised with new data, our eight individual models explained 8–73% of the variance in betacoronavirus positivity (mean of 32%), with the ensemble *R*^*2*^ increasing to 60% (Supplemental Figure 2). Using our previously applied 90% sensitivity threshold, our revised ensemble identified a more narrow set of 311 bat species as likely undetected hosts of betacoronaviruses. Predictions from the initial and revised ensembles were strongly correlated (ρ = 0.97). However, after dynamically updating our models, our revised ensemble lost 54 suspect reservoirs and gained 26 new suspect reservoirs (Figure 4A). Predicted reservoir species that were lost from the initial ensemble were dominated by members of the Vespertillionidae, whereas new suspect hosts were gained in the Vespertillionidae, Hipposideridae, and Molossidae.

**Figure 4.**
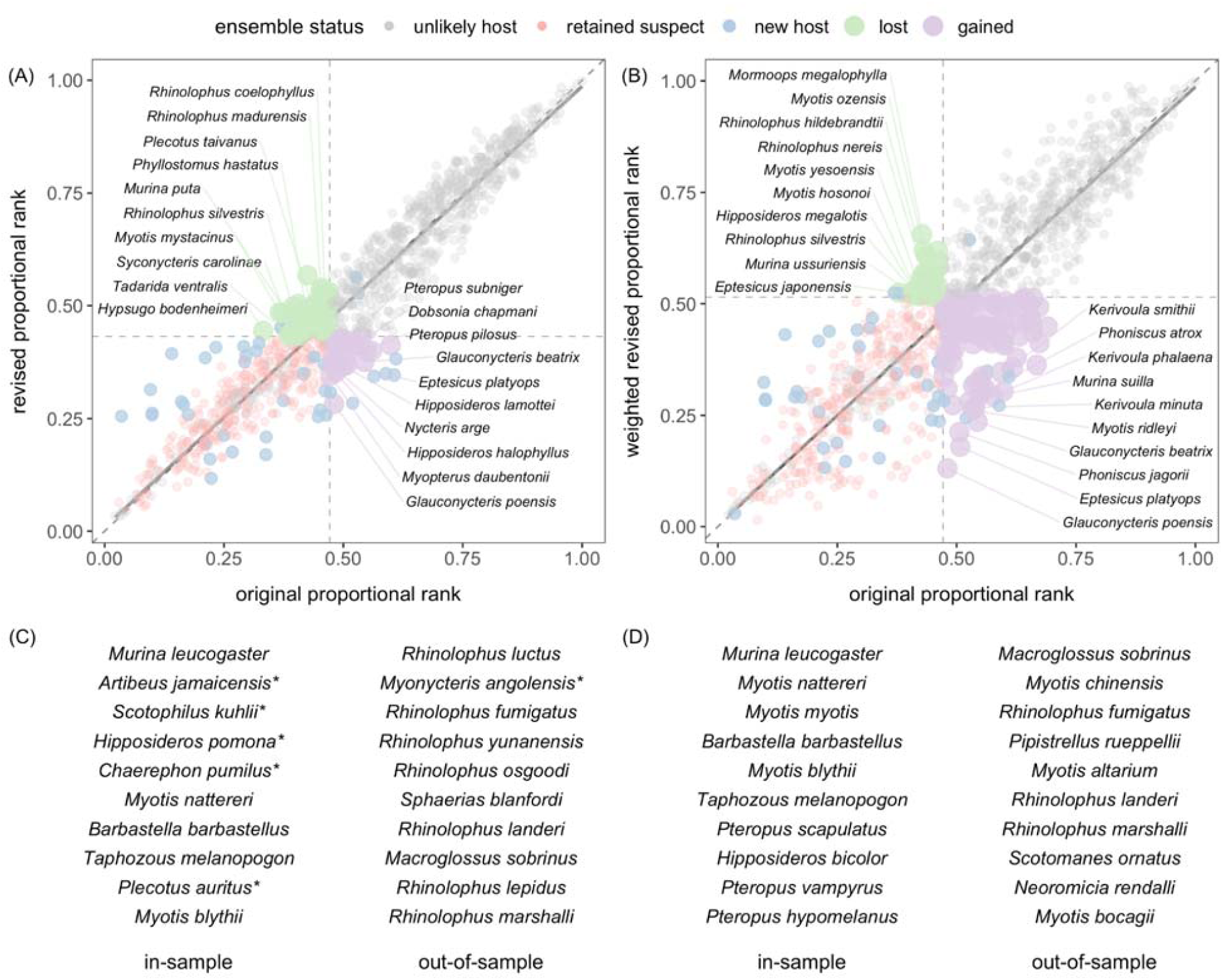
Comparing bat betacoronavirus host prediction with dynamic model updates. Scatterplots show bat species predictions from our original ensemble in 2020 against (A) the revised predictions after updating models with 40 new hosts and (B) the final predictions from the weighted revised ensemble.Species are colored by their status in the respective revised ensemble. Trendlines show a linear regression fit between original and revised predictions against a 1:1 line, whereas dashed lines display the threshold cutoffs from each ensemble. The top 10 in- and out-of-sample predictions from the original (C) and final (D) ensemble. Asterisks indicate that five of the original top 10 in-sample predictions, and one of the top 10 out-of-sample predictions, have been empirically confirmed since the first iteration of our study.

Using the 40 newly discovered hosts, we were also able to tailor the updated ensemble responsively to model performance. To do so, we weighted the rank averaging across models based on their AUC-TPTSC score relative to the lowest performing model (Network-2). In doing so, we effectively dropped Network-2 from the ensemble, a choice supported by the fact the model’s predictions are substantially poorer than expected at random. In the updated ensemble, this correction creates a marginal improvement in model performance (unweighted ensemble: AUC-TPTSC = 0.755; weighted ensemble: AUC-TPTSC = 0.788). Therefore, we applied this weighting to the ensemble of updated predictions in the final copy released with this study.

This weighted revised ensemble identified 423 suspect bat hosts, dramatically expanding the scope of plausible candidates for future virus discovery compared to the two prior unweighted ensembles (Figure 4B). Predictions from this final ensemble iteration were slightly less correlated with those from our initial ensemble (ρ = 0.92), and these final predictions retained most suspect hosts from the original ensemble. The top-ranked undiscovered hosts retained between our model updates included *Murina leucogaster, Myotis nattereri, M. blythii, Barbastella barbastellus*, and *Taphozous melanopogon* in-sample, whereas the top out-of-sample hosts consistent between ensembles included *Macroglossus sobrinus, Rhinolophus fumigatus*, and *R. marshalli* (Figure 4C). Only 26 predicted hosts were lost, most of which were from the Vespertilionidae and *Rhinolophus* genus. Of the 208 additional predicted hosts added to our final ensemble, most of these bat species were observed in the families Vespertilionidae (primarily the genus *Myotis*), Pteropodidae (primarily the genus *Pteropus*), Molossidae (primarily the genera *Chaerephon* and *Mops*), Hipposideridae (all in the genus *Hipposideros*), and Rhinolophidae, although we also identified several new predicted betacoronavirus hosts in the families Nycteridae, Phyllostomidae, Emballonuridae, and Mystacinidae. The top-ranked novel predicted in-sample hosts included *Myotis myotis, Hipposideros bicolor*, and multiple *Pteropus* species (*P. scapulatus, P. vampyrus*, and *P. hypomelanus*), whereas the most likely new out-of-sample hosts included multiple *Myotis* species (*M. chinensis, M. altarium*, and *M. bocagii*), *Pipistrellus rueppellii, Scotomanes ornatus, Neoromicia rendalli*, and *Rhinolophus landeri* (Figure 4D).

For *Rhinolophus* bats specifically, our final ensemble identified 11 new suspect hosts, resulting in likely hosts for over three quarters of the species in this genus that are not currently known to be infected by betacoronaviruses (44 of 57 in our dataset), compared to 20 known hosts. Given the known roles of rhinolophid bats as hosts of SARS-like CoVs^18,21,50^, it is notable that our results suggest the diversity of these viruses could be undescribed for around three-fourths of these potential reservoir bat species.

As in our initial ensemble, we lastly evaluated geographic and taxonomic patterns in this finalized set of predicted betacoronavirus hosts. Spatially, undiscovered bat hosts were globally distributed (in not only the eastern but also western hemisphere) and especially concentrated within a more narrow band of equatorial sub-Saharan Africa and more starkly in Malaysia and Borneo (Figure 5A). Importantly, the geography of these predicted hosts contrasted with the distributions of both known bat hosts and likely hosts from our initial ensemble, each of which instead showed a stronger hotspot in southern China. We also identified distinct clades of high-risk bat hosts from the weighted revised ensemble (Figure 5B, Supplemental Table 2). Both the *Rhinolophus* genus and subclades of the Pteropodidae again had greater concentrations of predicted betacoronavirus hosts, although phylogenetic factorization now identified the Old World molossids (i.e., genus *Mops and Chaerophon*) as particularly likely to host these viruses, even though the Mollosidae as a whole have lower mean probabilities of betacoronavirus hosts.

**Figure 5.**
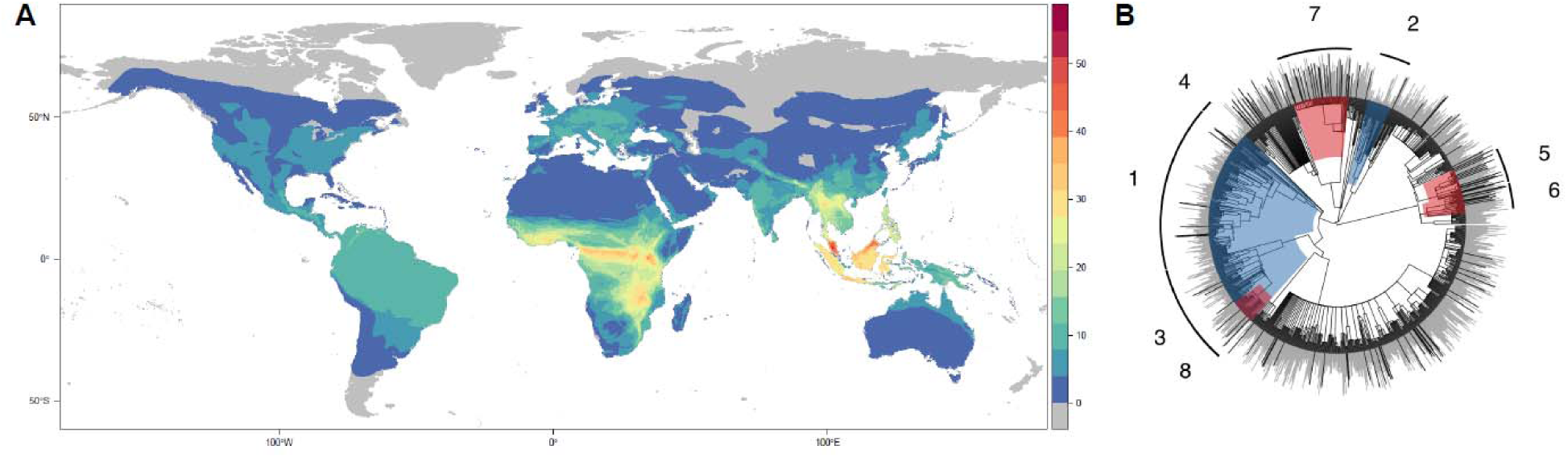
Predicted geographic and evolutionary hotspots of bat betacoronavirus hosts (updated ensemble predictions). (A) In the weighted revised ensemble predictions, most predicted undiscovered betacoronavirus hosts are found in sub-Saharan Africa and southeast Asia, especially in Malaysia and Borneo (and less in the high-elevation mainland hotspot where most reservoirs of SARS-like viruses are currently known). Predicted hosts from this final ensemble were also most likely in the *Rhinolophus* genus (clade 7), several subclades of the Pteropodidae (clades 5 and 6), and the Old World molossids (clade 8), even though the Mollosidae as a whole had less likely hosts (clade 3). Bar height in th phylogeny indicates predicted rank, and colors indicate clades identified through phylogenetic factorization (red indicates clades more likely to contain hosts, blue indicates less likely hosts; see Supplemental Table 2 for more information).

These geographic hotspots and clade-specific patterns of predictions could be particularly applicable for guiding future viral discovery and surveillance. On the one hand, betacoronavirus sampling in southeast Asian bat taxa (especially the genus *Rhinolophus*) may have a high success of viral detection (and isolation) of sarbecoviruses specifically but may not substantially improve existing bat sampling gaps^5^. On the other hand, discovery of novel betacoronaviruses in high-priority pteropodid clades, Old World molossids, and Neotropical bats could significantly revise our understanding of the bat–virus association network relative to the coevolutionary distribution of bat betacoronaviruses^38^. For example, predicted bat hosts in the Neotropics may be unlikely reservoirs of sarbecoviruses (given their known distribution in the eastern hemisphere) but would be expected to carry novel viruses from the subgenus *Merbecovirus*. Such discoveries could be particularly important for global health security, given the surprising recent identification of MERS-like viruses within the merbecoviruses in Mexican and Belizean bats^31,58^ and the likelihood that post-COVID research efforts will focus disproportionately on Asia despite the near-global presence of bat betacoronavirus hosts.

### Insights into SARS-CoV-2’s emergence

Our work suggests that over 400 bats may host undiscovered betacoronaviruses and that these species can be prioritized for sampling more efficiently via machine learning. Although our models do not target sarbecoviruses specifically, these efforts may help find more SARS-like viruses in wildlife and may even uncover the direct progenitor of SARS-CoV-2, particularly given that 44 species of horseshoe bats are predicted to host undiscovered betacoronaviruses. However, our models provide otherwise limited insights into the origins of SARS-CoV-2, given the likely role of non-bat bridge hosts in spillover to humans^59,60^. We thus attempted a similar model ensemble in June 2020, using five of our eight models to predict the broader mammal–virus network with a focus on potential betacoronavirus bridge hosts. At the time, only 30 non-bat hosts of betacoronaviruses were available in our data. Among the five models, we found poor concordance in predictions (Supplemental Figure 3). Predictions were also heavily biased towards well-studied and domesticated mammals (e.g., *Ovis aries, Vulpes vulpes, Capra hircus, Procyon lotor, Rattus rattus*), indicating that sampling bias dwarfed biological signals. As such, we evaluated these models as having limited value or consistency for an ensemble. This finding may be relevant given other studies have also modeled susceptibility to SARS-CoV-2 across mammals; however, some have used more detailed trait data and thus likely make better predictions at this broader taxonomic scale^61,62^

Instead of further calibrating this mammal-wide ensemble, we focused on the outputs of Trait-3, which predicts how species should share viruses in nature based on their evolutionary history and geography. In June 2020, we predicted the mammals expected to share viruses with *Rhinolophus affinis* and *R. malayanus*, which hosted the two viruses (RaTG13 and RmYN02) most relevant to SARS-CoV-2’s origins known at that time^23,63^ (a closer related virus, RpYN06, has since been discovered in *R. pusillus*^*64*^). We predicted that these two bat species are disproportionately likely to share viruses with pangolins (Pholidota) and carnivores (Carnivora), including civets (Viverridae), mustelids (Mustelidae), and cats (Felidae; Supplemental Figure 4). These predictions have been broadly validated by the role of the masked palm civet (*Paguma larvata*) in the original SARS-CoV outbreak^65,66^, the discovery of SARS-CoV-2-like viruses in the Sunda pangolin (*Manis javanica*)^34^, and extensive “spillback” of SARS-CoV-2 into captive big cats, domestic cats, and both farmed and wild mink^67,68^. Notably, only the association between palm civets and SARS-CoV was present in the training data used for generating predictions from Trait-3.

Given these successful predictions, we expect there might be potential insights into SARS-CoV-2’s emergence when these predictions are paired with data on wildlife supply chains. Out of the top 30 species (Figure 6), two are known to have been traded in wildlife markets in Wuhan immediately before the pandemic (the wild boar, *Sus scrofa*; the palm civet, *Paguma larvata*), as were two species closely related to those in the top predictions (the Siberian weasel, *Mustela siberica*; the Northern hog badger, *Arctonyx albogularis*)^69^. Another top species, the Burmese ferret badger (*Melogale personata*), was also reportedly of interest in the World Health Organization’s origins investigation^70^. Our models indicate that any of these species would be expected to regularly share viruses with relevant *Rhinolophus* bats in nature. Although bats use habitats differently than most of these likely bridge host species, opportunities for contact exist: a recent study from Gabon found cohabitation among pangolins, bats, and other mammals in burrows^71^. A number of species potentially implicated in the origins of SARS-CoV-2 could therefore have plausibly acquired a progenitor to SARS-CoV-2 in nature, at some point prior to contact with—and spillover into—humans (but see^72^). We suggest that this shortlist of species (Figure 6) may therefore be a useful line of evidence for further investigations into the identity of potential bridge hosts, especially in combination with experimental evaluation of susceptibility.

**Figure 6.**
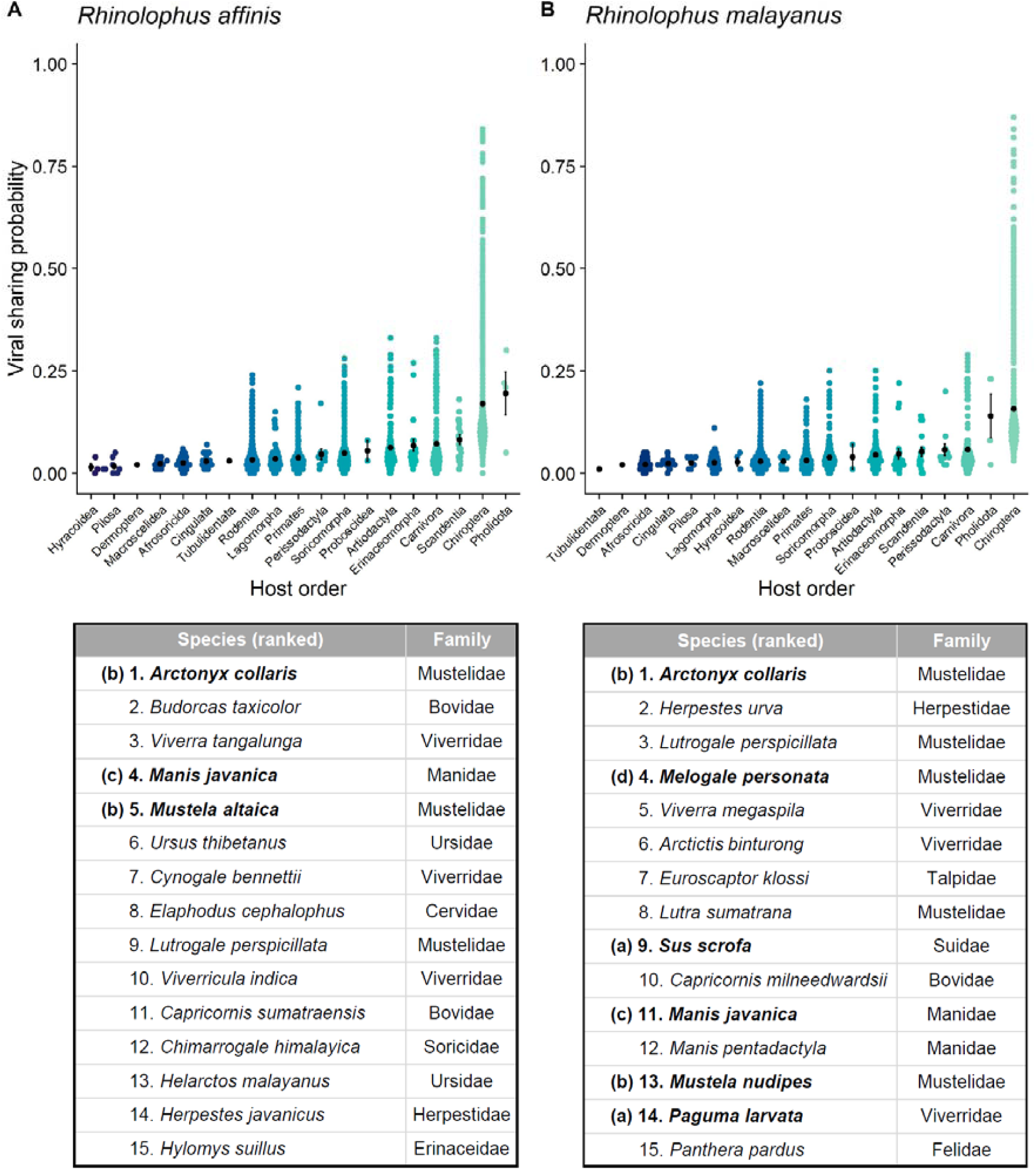
Potential bridge hosts involved in SARS-CoV-2’s emergence. Each dot represents predicted species-level sharing probabilities with (A) *Rhinolophus affinis* and (B) *Rhinolophus malayanus*, estimated according to the phylogeographic viral sharing model (Trait-3)^101^. Each coloured point is a mammal species. (Black points and error bars denote means and standard errors for each order; mammal orders are arranged according to their mean sharing probability.) Tables below report the top 15 predicted non-bat species for each; several families are disproportionately represented, including pangolins (Pholidota: Manidae), mustelids (Carnivora: Mustelidae),and civets (Carnivora: Viverridae). Notable species are bolded: (a) the wild boar *S. scrofa* and palm civet *P. larvata* were both traded in wildlife markets in Wuhan prior to the pandemic, as were (b) close relatives of the Greater hog badger, *A. collaris* (the Northern hog badger, *A. albogularis*), and of the mountain weasel, *M. altaica*, and Malayan weasel, *M. nudipes* (the Siberian weasel, *M. siberica*). (c) SARS-CoV-2-like viruses have been found in traded Sunda pangolins (*M. javanica*) outside of Wuhan, though the species was not reported in Wuhan. (d) The ferret badger (*M. personata*) was also reportedly of interest in the World Health Organization’s origins investigation, which explored the role of wildlife farm supply chains.

## Conclusions

Our study is the first to demonstrate through predictive validation that machine learning models could help optimize wildlife sampling for undiscovered viruses. As such, the growing toolkit of models that predict host–pathogen interactions are likely to aid future efforts both to predict and prevent pandemics and to trace the origins of novel infections after they emerge. However, these tools will work best if they are implemented through a dynamic process of prediction, data collection, validation, and updating, as we have implemented in this study. Although some studies have incidentally tested specific hypotheses (e.g., filovirus models and bat surveys^7,11^, henipavirus models and experimental infections^2,16^, vector–virus models and competence trials^13–15^), predictions are almost never subject to systematic verification. Greater dialogue between modelers and empiricists is necessary to confront this research gap. This is particularly important when establishing species’ role as viral reservoirs rather than incidental hosts; susceptibility is only one aspect of host competence^1,10^. Future work, including longitudinal tracking of viral shedding over space and time, isolation of live virus from wild animals, and experimental confirmation of viral replication, can support more robust conclusions about whether predicted host species actually play a role in viral maintenance^16^ as well as inform related efforts to pinpoint and minimize risk factors for pathogen spillover^73,74^.

Our study is also the first to benchmark the performance of a set of differently calibrated and designed statistical models all trained for one host prediction task. We found a surprising range of model performance, even among a set of models that all performed well on training data. This finding underscores the need to incorporate long-term validation into similar studies and suggests there may be key lessons about viral ecology to be learned from this kind of exercise. In our study, we found that network models performed mostly at random in validation against new bat hosts, whereas trait-based models were more successful in their predictions. Two explanations may underscore this difference in model performance. First, models that successfully predict the broader mammal–virus network are likely to vary in performance when subset to any given node. This may seem contradictory to the idea that understanding the broader “rules of life” underpinning mammal–virus interactions will improve predictions in specific cases. However, there are ways to combine the strengths of both network-and trait-based approaches. In a closely related study, network predictions of zoonotic risk performed essentially at random, whereas a hybrid approach that embedded network predictions in targeted, trait-based models performed better than any approach in isolation^75^. Future work should aim to develop and benchmark these kinds of hybrid model approaches more extensively and treat exclusively network-driven predictions with caution in the interim.

Second, models that integrate data on host ecology, evolution, and biogeography are likely to make more powerful predictions than those that are mostly agnostic to biology. This has many broader implications. Most importantly, it suggests that filling gaps in the basic biology of bats is a key step towards zoonotic risk assessment and can benefit both pandemic prevention and bat conservation. In particular, high-quality host genomes are critical to developing better predictive features, including genome composition bias metrics, improved host phylogenetic trees, and immunological traits^62,76–79^. Whole-genome sequencing through initiatives such as the Bat1K Project (https://bat1k.ucd.ie) will expand the sparsely available data on bat genomics and can facilitate other insights into the immune pathways used by bats to harbor virulent viruses^80–82^. Targeted sequencing could also identify endogenized viral elements in bat genomes, shedding light on bat virus diversity and the evolution of bat immune systems^83,84^. Large-scale research networks, such as the Global Union of Bat Diversity Networks (https://gbatnet.blogspot.com) and its member networks, will further facilitate efficient sample sharing and ensure proper partnerships and equitable access and benefit sharing of knowledge across countries^85,86^. Additionally, museum specimens and historical collections offer important opportunities to retrospectively screen samples for betacoronaviruses (thereby testing predictions), sequence tissue for assembly of host genomes, and enhance our understanding of complex host–virus interactions^87^.

Lastly, our iterative modeling of bat betacoronaviruses fits into a broader set of synergies in One Health research on bats, which can create win–win scenarios for conservation and outbreak prevention. For example, North American bats are threatened by an emerging disease, white-nose syndrome^88^, which has documented synzootic interactions with other bat coronaviruses^89^; at least seven North American bat species that can be infected by the fungal pathogen (*Eptesicus fuscus, Myotis ciliolabrum, M. lucifugus, M. septentrionalis, M. velifer, M. volans*, and *Tadarida brasiliensis*) are among the 423 bat species we predict could be undiscovered betacoronavirus hosts. Although our predictions do not imply bat susceptibility to SARS-CoV-2 specifically (and experimental infections of *E. fuscus* have been unsuccessful^90^), efforts to minimize risks of SARS-CoV-2 “spillback” into novel bat reservoirs^91–95^, as well as to understand the dynamics of other bat coronaviruses, will both reduce zoonotic risk and help understand and counteract disease-related population declines. Similarly, conservationists have expressed concern that negative framing of bats as the source of SARS-CoV-2 has impacted public and governmental attitudes toward bat conservation^96^; this can fuel negative responses, including indiscriminate culling (i.e., reduction of populations by slaughter), which has already occured in response to COVID-19 even outside of Asia (where spillover likely occurred)^97^. Evidence demonstrates that culling has numerous negative consequences, not only threatening population viability^98^ but also possibly increasing viral transmission within the very species that are targeted^99,100^. Bat conservation programs and One Health practitioners must continue to work together to find sustainable solutions for humans to live safely alongside wildlife and to communicate with the public about the ecological importance of these highly vulnerable species and the science of pathogen spillover.

## Acknowledgements

We thank Heather Wells for generously sharing thoughtful comments and code. The Viral Emergence Research Initiative (VERENA) consortium is supported by L’Institut de Valorisation de Données (IVADO) through Université de Montreal and by NSF BII 2021909; for more information, see viralemergence.org. LMB was supported by the Wellcome Trust (217221/Z/19/Z). MS was supported by the Research Foundation -Flanders (FWO17/PDO/067) and the Flemish Government under the “Onderzoeksprogramma Artificiële Intelligentie (AI) Vlaanderen” program. Lastly, we thank three anonymous reviewers for constructive feedback.

## Data and Code Availability

The standardized data on betacoronavirus associations, and all associated predictor data, is available from the VERENA consortium’s Github (github.com/viralemergence/virionette). All modeling teams contributed an individual repository with their methods, which are available in the organizational directory (github.com/viralemergence). Code for the 2020 analysis, and a working reproduction of each author’s contributions, is available from the study repository (github.com/viralemergence/Fresnel); code to produce the updated analyses from 2021 is available in a separate repository (github.com/viralemergence/Fresnel_Jun). A complete list of the 423 predicted betacoronavirus bat hosts from our final ensemble is provided in Supplemental File 1.

## Bibliography

1 Viana M, Mancy R, Biek R, et al. Assembling evidence for identifying reservoirs of infection. Trends Ecol Evol 2014; 29: 270–9.

2 Plowright RK, Becker DJ, Crowley DE, et al. Prioritizing surveillance of Nipah virus in India. PLoS Negl Trop Dis 2019; 13: e0007393.

3 Becker DJ, Crowley DE, Washburne AD, Plowright RK. Temporal and spatial limitations in global surveillance for bat filoviruses and henipaviruses. Biol Lett 2019; 15: 20190423.

4 Washburne AD, Crowley DE, Becker DJ, et al. Taxonomic patterns in the zoonotic potential of mammalian viruses. PeerJ 2018; 6: e5979.

5 Crowley D, Becker D, Washburne A, Plowright R. Identifying Suspect Bat Reservoirs of Emerging Infections. Vaccines. 2020; 8: 228.

6 Becker DJ, Washburne AD, Faust CL, Mordecai EA, Plowright RK. The problem of scale in the prediction and management of pathogen spillover. Philos Trans R Soc Lond B Biol Sci 2019; 374: 20190224.

7 Han BA, Schmidt JP, Alexander LW, Bowden SE, Hayman DTS, Drake JM. Undiscovered Bat Hosts of Filoviruses. PLoS Negl Trop Dis 2016; 10: e0004815.

8 Han BA, Majumdar S, Calmon FP, et al. Confronting data sparsity to identify potential sources of Zika virus spillover infection among primates. Epidemics 2019; 27: 59–65.

9 Becker DJ, Han BA. The macroecology and evolution of avian competence for Borrelia burgdorferi. DOI:10.1101/2020.04.15.040352.

10 Becker DJ, Seifert SN, Carlson CJ. Beyond Infection: Integrating Competence into Reservoir Host Prediction. Trends Ecol Evol 2020; published online Sept 11. DOI:10.1016/j.tree.2020.08.014.

11 Yang X-L, Zhang Y-Z, Jiang R-D, et al. Genetically Diverse Filoviruses in Rousettus and Eonycteris spp. Bats, China, 2009 and 2015. Emerg Infect Dis 2017; 23: 482–6.

12 Laing ED, Mendenhall IH, Linster M, et al. Serologic Evidence of Fruit Bat Exposure to Filoviruses, Singapore, 2011-2016. Emerg Infect Dis 2018; 24: 114–7.

13 Evans MV, Dallas TA, Han BA, Murdock CC, Drake JM. Data-driven identification of potential Zika virus vectors. Elife 2017; 6. DOI:10.7554/eLife.22053.

14 Guedes DR, Paiva MH, Donato MM, et al. Zika virus replication in the mosquito Culex quinquefasciatus in Brazil. Emerg Microbes Infect 2017; 6: e69.

15 Smartt CT, Shin D, Kang S, Tabachnick WJ. Culex quinquefasciatus (Diptera: Culicidae) From Florida Transmitted Zika Virus. Front Microbiol 2018; 9: 768.

16 Seifert SN, Letko MC, Bushmaker T, et al. Rousettus aegyptiacus Bats Do Not Support Productive Nipah Virus Replication. The Journal of Infectious Diseases. 2020; 221: S407–13.

17 Gokhale, Sreelekshmy M, Sudeep AB, et al. Detection of possible Nipah virus infection in Rousettus leschenaultii and Pipistrellus pipistrellus bats in Maharashtra, India. J Infect Public Health 2021; published online June 11. DOI:10.1016/j.jiph.2021.05.001.

18 Anthony SJ, Johnson CK, Greig DJ, et al. Global patterns in coronavirus diversity. Virus Evol 2017; 3: vex012.

19 Denison MR, Graham RL, Donaldson EF, Eckerle LD, Baric RS. Coronaviruses: an RNA proofreading machine regulates replication fidelity and diversity. RNA Biol 2011; 8: 270–9.

20 Ren W, Li W, Yu M, et al. Full-length genome sequences of two SARS-like coronaviruses in horseshoe bats and genetic variation analysis. J Gen Virol 2006; 87: 3355–9.

21 Li W, Shi Z, Yu M, et al. Bats are natural reservoirs of SARS-like coronaviruses. Science 2005; 310: 676–9.

22 Yang X-L, Hu B, Wang B, et al. Isolation and Characterization of a Novel Bat Coronavirus Closely Related to the Direct Progenitor of Severe Acute Respiratory Syndrome Coronavirus. J Virol 2015; 90: 3253–6.

23 Zhou P, Yang X-L, Wang X-G, et al. A pneumonia outbreak associated with a new coronavirus of probable bat origin. Nature 2020; 579: 270–3.

24 Guan Y, Zheng BJ, He YQ, et al. Isolation and characterization of viruses related to the SARS coronavirus from animals in southern China. Science 2003; 302: 276–8.

25 Hu B, Zeng L-P, Yang X-L, et al. Discovery of a rich gene pool of bat SARS-related coronaviruses provides new insights into the origin of SARS coronavirus. PLoS Pathog 2017; 13: e1006698.

26 Memish ZA, Mishra N, Olival KJ, et al. Middle East respiratory syndrome coronavirus in bats, Saudi Arabia. Emerg Infect Dis 2013; 19: 1819–23.

27 Wang Q, Qi J, Yuan Y, et al. Bat origins of MERS-CoV supported by bat coronavirus HKU4 usage of human receptor CD26. Cell Host Microbe 2014; 16: 328–37.

28 Yang Y, Du L, Liu C, et al. Receptor usage and cell entry of bat coronavirus HKU4 provide insight into bat-to-human transmission of MERS coronavirus. Proc Natl Acad Sci U S A 2014; 111: 12516– 21.

29 Hu B, Ge X, Wang L-F, Shi Z. Bat origin of human coronaviruses. Virology Journal. 2015; 12. DOI:10.1186/s12985-015-0422-1.

30 Anthony SJ, Gilardi K, Menachery VD, et al. Further Evidence for Bats as the Evolutionary Source of Middle East Respiratory Syndrome Coronavirus. MBio 2017; 8. DOI:10.1128/mBio.00373-17.

31 Anthony SJ, Ojeda-Flores R, Rico-Chávez O, et al. Coronaviruses in bats from Mexico. J Gen Virol 2013; 94: 1028–38.

32 Yang L, Wu Z, Ren X, et al. MERS-related betacoronavirus in Vespertilio superans bats, China. Emerg Infect Dis 2014; 20: 1260–2.

33 Nielsen R, Wang H, Pipes L. Synonymous mutations and the molecular evolution of SARS-Cov-2 origins. DOI:10.1101/2020.04.20.052019.

34 Lam TT-Y, Shum MH-H, Zhu H-C, et al. Identifying SARS-CoV-2 related coronaviruses in Malayan pangolins. Nature 2020; published online March 26. DOI:10.1038/s41586-020-2169-0.

35 Xiao K, Zhai J, Feng Y, et al. Isolation of SARS-CoV-2-related coronavirus from Malayan pangolins. Nature 2020; published online May 7. DOI:10.1038/s41586-020-2313-x.

36 Zhang T, Wu Q, Zhang Z. Probable Pangolin Origin of SARS-CoV-2 Associated with the COVID-19 Outbreak. Curr Biol 2020; 30: 1578.

37 Andersen KG, Rambaut A, Ian Lipkin W, Holmes EC, Garry RF. The proximal origin of SARS-CoV-2. Nature Medicine. 2020; 26: 450–2.

38 Olival KJ, Hosseini PR, Zambrana-Torrelio C, Ross N, Bogich TL, Daszak P. Host and viral traits predict zoonotic spillover from mammals. Nature 2017; 546: 646–50.

39 Fritz SA, Bininda-Emonds ORP, Purvis A. Geographical variation in predictors of mammalian extinction risk: big is bad, but only in the tropics. Ecol Lett 2009; 12: 538–49.

40 Jones KE, Bielby J, Cardillo M, et al. PanTHERIA: a species-level database of life history, ecology, and geography of extant and recently extinct mammals. Ecology. 2009; 90: 2648–2648.

41 Wilman H, Belmaker J, Simpson J, de la Rosa C, Rivadeneira MM, Jetz W. EltonTraits 1.0: Species-level foraging attributes of the world’s birds and mammals. Ecology. 2014; 95: 2027–2027.

42 Trifonova N, Kenny A, Maxwell D, Duplisea D, Fernandes J, Tucker A. Spatio-temporal Bayesian network models with latent variables for revealing trophic dynamics and functional networks in fisheries ecology. Ecological Informatics. 2015; 30: 142–58.

43 Rohr RP, Scherer H, Kehrli P, Mazza C, Bersier L-F. Modeling food webs: exploring unexplained structure using latent traits. Am Nat 2010; 176: 170–7.

44 Dallas T, Park AW, Drake JM. Predicting cryptic links in host-parasite networks. PLoS Comput Biol 2017; 13: e1005557.

45 Han BA, Schmidt JP, Bowden SE, Drake JM. Rodent reservoirs of future zoonotic diseases. Proceedings of the National Academy of Sciences. 2015; 112: 7039–44.

46 Washburne AD, Silverman JD, Morton JT, et al. Phylofactorization: a graph partitioning algorithm to identify phylogenetic scales of ecological data. Ecol Monogr 2019; 89: e01353.

47 Brandão PE, Scheffer K, Villarreal LY, et al. A coronavirus detected in the vampire bat Desmodus rotundus. Braz J Infect Dis 2008; 12: 466–8.

48 Corman VM, Rasche A, Diallo TD, et al. Highly diversified coronaviruses in neotropical bats. J Gen Virol 2013; 94: 1984–94.

49 Moreira-Soto A, Taylor-Castillo L, Vargas-Vargas N, Rodríguez-Herrera B, Jimenez C, Corrales-Aguilar E. Neotropical bats from Costa Rica harbour diverse coronaviruses. Zoonoses Public Health 2015; 62: 501–5.

50 Wang L, Fu S, Cao Y, et al. Discovery and genetic analysis of novel coronaviruses in least horseshoe bats in southwestern China. Emerg Microbes Infect 2017; 6: e14.

51 Lin X-D, Wang W, Hao Z-Y, et al. Extensive diversity of coronaviruses in bats from China. Virology. 2017; 507: 1–10.

52 Wacharapluesadee S, Duengkae P, Rodpan A, et al. Diversity of coronavirus in bats from Eastern Thailand. Virol J 2015; 12: 57.

53 Guy C, Ratcliffe JM, Mideo N. The influence of bat ecology on viral diversity and reservoir status. Ecol Evol 2020; 2008: 209.

54 Wacharapluesadee S, Duengkae P, Chaiyes A, et al. Longitudinal study of age-specific pattern of coronavirus infection in Lyle’s flying fox (Pteropus lylei) in Thailand. Virol J 2018; 15: 38.

55 Bergner LM, Orton RJ, Broos A, et al. Diversification of mammalian deltaviruses by host shifting. 2020; : 2020.06.17.156745.

56 Bergner LM, Orton RJ, da Silva Filipe A, et al. Using noninvasive metagenomics to characterize viral communities from wildlife. Mol Ecol Resour 2019; 19: 128–43.

57 Bergner LM, Orton RJ, Streicker DG. Complete Genome Sequence of an Alphacoronavirus from Common Vampire Bats in Peru. Microbiol Resour Announc 2020; 9. DOI:10.1128/MRA.00742-20.

58 Neely BA, Janech MG, Brock Fenton M, Simmons NB, Bland AM, Becker DJ. Surveying the vampire bat (Desmodus rotundus) serum proteome: a resource for identifying immunological proteins and detecting pathogens. Cold Spring Harbor Laboratory. 2020; : 2020.12.04.411660.

59 Zhao J, Cui W, Tian B-P. The Potential Intermediate Hosts for SARS-CoV-2. Frontiers in Microbiology. 2020; 11. DOI:10.3389/fmicb.2020.580137.

60 Cui J, Li F, Shi Z-L. Origin and evolution of pathogenic coronaviruses. Nature Reviews Microbiology. 2019; 17: 181–92.

61 Fischhoff IR, Castellanos AA, Rodrigues JPGLM, Varsani A, Han BA. Predicting the zoonotic capacity of mammal species for SARS-CoV-2. bioRxiv 2021; published online Feb 19. DOI:10.1101/2021.02.18.431844.

62 Wardeh M, Baylis M, Blagrove MSC. Predicting mammalian hosts in which novel coronaviruses can be generated. Nat Commun 2021; 12: 780.

63 Zhou H, Chen X, Hu T, et al. A novel bat coronavirus reveals natural insertions at the S1/S2 cleavage site of the Spike protein and a possible recombinant origin of HCoV-19. DOI:10.1101/2020.03.02.974139.

64 Zhou H, Ji J, Chen X, et al. Identification of novel bat coronaviruses sheds light on the evolutionary origins of SARS-CoV-2 and related viruses. Cell 2021; published online June 9. DOI:10.1016/j.cell.2021.06.008.

65 Wang M, Yan M, Xu H, et al. SARS-CoV Infection in a Restaurant from Palm Civet. Emerging Infectious Diseases. 2005; 11: 1860–5.

66 Song H-D, Tu C-C, Zhang G-W, et al. Cross-host evolution of severe acute respiratory syndrome coronavirus in palm civet and human. Proc Natl Acad Sci U S A 2005; 102: 2430–5.

67 Jia P, Dai S, Wu T, Yang S. New Approaches to Anticipate the Risk of Reverse Zoonosis. Trends Ecol Evol 2021; 36: 580–90.

68 Fagre AC, Cohen L, Eskew EA, et al. Spillback in the Anthropocene: the risk of human-to-wildlife pathogen transmission for conservation and public health. DOI:10.32942/osf.io/sx6p8.

69 Xiao X, Newman C, Buesching CD, Macdonald DW, Zhou Z-M. Animal sales from Wuhan wet markets immediately prior to the COVID-19 pandemic. Sci Rep 2021; 11: 11898.

70 McKay B, Page J, Hinshaw D. In Hunt for Covid-19 Origin, WHO Team Focuses on Two Animal Types in China. Wall Street Journal. 2021; published online Feb 18.

71 Lehmann D, Halbwax ML, Makaga L, et al. Pangolins and bats living together in underground burrows in Lopé National Park, Gabon. African Journal of Ecology. 2020. DOI:10.1111/aje.12759.

72 Lee J, Hughes T, Lee M-H, et al. No Evidence of Coronaviruses or Other Potentially Zoonotic Viruses in Sunda pangolins (Manis javanica) Entering the Wildlife Trade via Malaysia. Ecohealth 2020; 17: 406–18.

73 Plowright RK, Becker DJ, McCallum H, Manlove KR. Sampling to elucidate the dynamics of infections in reservoir hosts. Philos Trans R Soc Lond B Biol Sci 2019; 374: 20180336.

74 Sokolow SH, Nova N, Pepin KM, et al. Ecological interventions to prevent and manage zoonotic pathogen spillover. Philos Trans R Soc Lond B Biol Sci 2019; 374: 20180342.

75 Poisot T, Ouellet M-A, Mollentze N, et al. Imputing the mammalian virome with linear filtering and singular value decomposition. arXiv [q-bio.QM]. 2021. http://arxiv.org/abs/2105.14973.

76 Babayan SA, Orton RJ, Streicker DG. Predicting reservoir hosts and arthropod vectors from evolutionary signatures in RNA virus genomes. Science 2018; 362: 577–80.

77 Mollentze N, Babayan SA, Streicker DG. Identifying and prioritizing potential human-infecting viruses from their genome sequences. Cold Spring Harbor Laboratory. 2020; : 2020.11.12.379917.

78 Rannala B, Yang Z. Phylogenetic inference using whole genomes. Annu Rev Genomics Hum Genet 2008; 9: 217–31.

79 Pedersen AB, Babayan SA. Wild immunology. Mol Ecol 2011; 20: 872–80.

80 Teeling EC, Vernes SC, Dávalos LM, et al. Bat Biology, Genomes, and the Bat1K Project: To Generate Chromosome-Level Genomes for All Living Bat Species. Annu Rev Anim Biosci 2018; 6: 23–46.

81 Jebb D, Huang Z, Pippel M, et al. Six reference-quality genomes reveal evolution of bat adaptations. Nature 2020; 583: 578–84.

82 Wang L-F, Gamage AM, Chan WOY, Hiller M, Teeling EC. Decoding bat immunity: the need for a coordinated research approach. Nat Rev Immunol 2021; 21: 269–71.

83 Skirmuntt EC, Escalera-Zamudio M, Teeling EC, Smith A, Katzourakis A. The Potential Role of Endogenous Viral Elements in the Evolution of Bats as Reservoirs for Zoonotic Viruses. Annu Rev Virol 2020; published online May 20. DOI:10.1146/annurev-virology-092818-015613.

84 Jebb D, Huang Z, Pippel M, et al. Six new reference-quality bat genomes illuminate the molecular basis and evolution of bat adaptations. 2019. https://pure.mpg.de/pubman/faces/ViewItemOverviewPage.jsp?itemId=item_3180152.

85 Kingston T, Aguirre L, Armstrong K, et al. Networking networks for global bat conservation. In: Bats in the Anthropocene: Conservation of Bats in a Changing World. Springer, Cham, 2016: 539– 69.

86 Phelps KL, Hamel L, Alhmoud N, et al. Bat Research Networks and Viral Surveillance: Gaps and Opportunities in Western Asia. Viruses 2019; 11. DOI:10.3390/v11030240.

87 de Souza Cortez J.L. Dunnum A.W. Ferguson F.A. Anwarali Khan D.L. Paul D.M. Reeder N.B. Simmons B.M. Thiers C.W. Thompson NS. Upham M.P.M. Vanhove P.W. Webala M. Weksler R. Yanagihara P.S. Soltis. CJASABAJBCACBMB. Integrating biodiversity infrastructure into pathogen discovery and mitigation of epidemic infectious diseases. Bioscience 2020. DOI:biaa064.

88 Frick WF, Pollock JF, Hicks AC, et al. An emerging disease causes regional population collapse of a common North American bat species. Science 2010; 329: 679–82.

89 Davy CM, Donaldson ME, Subudhi S, et al. White-nose syndrome is associated with increased replication of a naturally persisting coronaviruses in bats. Sci Rep 2018; 8: 15508.

90 Hall JS, Knowles S, Nashold SW, et al. Experimental challenge of a North American bat species, big brown bat (Eptesicus fuscus), with SARS-CoV-2. Transbound Emerg Dis 2020; published online Dec 9. DOI:10.1111/tbed.13949.

91 Olival KJ, Cryan PM, Amman BR, et al. Possibility for reverse zoonotic transmission of SARS-CoV-2 to free-ranging wildlife: A case study of bats. PLoS Pathog 2020; 16: e1008758.

92 Cox-Witton K, Baker ML, Edson D, Peel AJ, Welbergen JA, Field H. Risk of SARS-CoV-2 transmission from humans to bats - An Australian assessment. One Health 2021; 13: 100247.

93 Common SM, Shadbolt T, Walsh K, Sainsbury AW. The risk from SARS□CoV□2 to bat species in england and mitigation options for conservation field workers. Transboundary and Emerging Diseases. 2021. DOI:10.1111/tbed.14035.

94 Cook JD, Grant EHC, Coleman JTH, Sleeman JM, Runge MC. Risks posed by SARS□CoV□2 to North American bats during winter fieldwork. Conservation Science and Practice. 2021; 3. DOI:10.1111/csp2.410.

95 Kingston T, Frick W, Kading R, et al. IUCN SSC Bat Specialist Group (BSG) Recommended Strategy for Researchers to Reduce the Risk of Transmission of SARS-CoV-2 from Humans to Bats (version 2.0). IUCN SSC Bat Specialist Group (BSG), 2021.

96 Zhao H. COVID-19 drives new threat to bats in China. Science 2020; 367: 1436.

97 MB Fenton, S Mubareka, SM Tsang, NB Simmons, DJ Becker. COVID-19 and threats to bats. FACETS 2020; : in press.

98 Aguiar LMS, Brito D, Machado RB. Do current vampire bat (Desmodus rotundus) population control practices pose a threat to Dekeyser’s nectar bat’s (Lonchophylla dekeyseri) long-term persistence in the Cerrado? Acta Chiropt 2010; 12: 275–82.

99 Streicker DG, Recuenco S, Valderrama W, et al. Ecological and anthropogenic drivers of rabies exposure in vampire bats: implications for transmission and control. Proc Biol Sci 2012; 279: 3384– 92.

100 Blackwood JC, Streicker DG, Altizer S, Rohani P. Resolving the roles of immunity, pathogenesis, and immigration for rabies persistence in vampire bats. Proc Natl Acad Sci U S A 2013; 110: 20837– 42.

101 Albery GF, Eskew EA, Ross N, Olival KJ. Predicting the global mammalian viral sharing network using phylogeography. Nat Commun 2020; 11: 2260.

