## Supporting Materials for "Optimizing predictive models to prioritize viral discovery in zoonotic reservoirs"

**Methods.**

The underlying conceptual aim of this study was to produce and synthesize several different models that predict and rank candidate reservoir host and bridge host species (each with different methods, assumptions, and framings), to synthesize these into a consensus list, and then to compare model performance on newly discovered betacoronavirus hosts. We broadly structured our study around two modeling targets: (1) produce rankings of likely bat hosts of betacoronaviruses and (2) identify potential non-bat mammal bridge hosts. We developed a novel synthesis of datasets that merged existing knowledge about the broader mammal–virus network with targeted data collection about coronaviruses; implemented eight modeling methods; synthesized these into an ensemble; and post-hoc identified taxonomic patterns in prediction using phylogenetic factorization. We then compiled data on new bat hosts of betacoronaviruses using a combination of GenBank scrapes, systematic literature searches, mining metagenomic and transcriptomic datasets, and newly released datasets. We then compared the performance of our models on identifying these newly discovered hosts and produced a finalized, revised list of candidate hosts for targeted virus discovery and virus surveillance.

**Host–Virus Association Data**

Entries were downloaded from GenBank on March 27 2020 using the following search terms: Coronavirus, Coronaviridae, Orthocoronavirinae Alphacoronavirus, Betacoronavirus, Gammacoronavirus, and Deltacoronavirus. Data were sorted using a Python script that saved all available metadata regarding accession number, division, submission date, entry title, organism, genus, genome length, host classification, country, collection date, PubMed ID, journal containing associated publication, publication year, genome completeness, and the gene sequenced. The dataset was cleaned to remove duplicate entries, using GenBank accession number, and entries that did not correspond to viral sequences, using GenBank division. After cleaning, 31,473 entries remained, of which 25,628 had metadata regarding host species.

Data from GenBank were merged with the Host-Pathogen Phylogeny Project (HP3) dataset^1^. The HP3 dataset consists of 2,805 associations between 754 mammal hosts and 586 virus species, compiled from the International Committee on Taxonomy of Viruses (ICTV) database, and manually cleaned over a period of five years. Data collection on HP3 began in 2010 and has been static since 2017, but it still represents one of the most complete datasets on the mammal virome published with a high standard of data documentation^2^. Several recent studies have used the HP3 dataset to produce statistical models of viral sharing or zoonotic potential^3–5^, making it a comparable reference for a multi-model ensemble study.

Because of naming inconsistencies both within GenBank and between the two datasets (HP3 and GenBank), we used a two-step pipeline for taxonomic reconciliation. Viral names were matched to the ICTV 2019 master species list, up to the sub-genus level. Host species names were matched against GBIF using their species API with an automated Julia script, and processed to a fully cleaned set of names. This led to an harmonized dataset representing a global list of mammal-virus associations, from which the bat-coronavirus data can be extracted for downstream and specific modeling efforts. Because the HP3 dataset used an older version of the ICTV master list, and because not all host names in the GenBank metadata could be matched by the GBIF species API (or could be solved unambiguously to the species level), some host-virus interactions were lost; this reinforces the need to careful data curation of taxonomic metadata if they are to enable and support predictive pipelines.

In total, our association dataset described a network of 1,731 associations between 710 host species and 72 virus genera, with *Betacoronavirus* hosts deliberately sampled at a higher level of completeness.

**Predictor Data**

*Phylogeny*

We used a supertree of extant mammals to unify modeling approaches incorporating host phylogeny^6^. Although more recent mammal supertrees exist, we used this particular phylogeny for consistency with trait datasets and several of the modeling frameworks included in our ensemble. We manually matched select bat species names between our edge list and this particular phylogeny. This included reverting any *Dermanura* to their former *Artibeus* designation (i.e., *D. phaeotis*, *D. cinerea*, *D. tolteca*)^7^, switching *Tadarida* species to either *Mops* or *Chaerophon* species (i.e., *Tadarida condylura* to *Mops condylurus*, *Tadarida plicata* to *Chaerephon plicatus*, *Tadarida pumila* to *Chaerephon pumilus*)^8^, and renaming *Myotis pilosus* to the more recent *Myotis ricketti*. *Chaerephon pusillus* was considered its own species but is now synonymous with *Chaerephon pumilus**^8^*. Minor discrepancies between virus data and our phylogeny were also corrected (*Hipposideros commersonii* to *Hipposideros commersoni* [although more recently changed to *Macronycteris commersoni*], *Rhinolophus hildebrandti* to *Rhinolophus hildebrandtii*, *Neoromicia nana* to *Neoromicia nanus*). In other cases, some recently revised genera in our edge list were modified to match former genera in the mammal supertree: *Parastrellus hesperus* to *Pipistrellus hesperus*, and *Perimyotis subflavus* to *Pipistrellus subflavus**^9^*. Lastly, some names in our edge list missing from the mammal supertree represent former subspecies being raised to full species rank, and names were reverted accordingly: *Artibeus* *planirostris* to *Artibeus jamaicensis*, *Miniopterus fuliginosus* to *Miniopterus schreibersii*, *Triaenops afer* to *Triaenops persicus*, and *Carollia sowelli* to *Carollia brevicauda*. Although we recognize that these are each now recognized as distinct species, in all cases our synonymized names are thought to be either sister taxa or very closely related.

*Ecological traits*

We used a previously published dataset of 63 ecological traits describing the morphology, life history, biogeography, and diet of 1,116 bat species. These data are drawn from a combination of PanTHERIA^10^, EltonTraits^11^, and the IUCN Red List range maps, and were previously cleaned in a study producing predictions of bat reservoirs of filoviruses^12^. Four redundant variables (two for human population density, mean potential evapotranspiration in range, and body mass) were eliminated prior to analyses, favoring variables with higher completeness.

*Correction for sampling bias*

To correct for sampling bias, in the style of several previous studies^1,3^, we used the number of peer-reviewed citations available on a given host as a measure of scientific sampling effort. We used the R package *easyPubMed* to scrape the number of citations in PubMed returned when searching each of the 1,116 bat names in the trait data on April 10, 2020.

**Initial Modeling Approaches**

We set out to develop an ensemble of models that could each use these data to predict bat hosts of betacoronaviruses. Many such algorithms exist, particularly given that a number of machine learning algorithms can all be applied to binary classification tasks. Our goal was not to benchmark slight differences between these algorithms, which other studies have explored more comprehensively, particularly in the species distribution modeling literature^13^. Rather, we aimed to sample a set of approaches that are broadly representative of the current state-of-the-art. Our team produced an initial ensemble of eight statistical models (Supplemental Table 1) and applied them to generate a predictive set of eight models for bats and five for other mammals. Four use a network-theoretic component (k-nearest neighbors, linear filtering, trait-free plug and play, and scaled-phylogeny), while three primarily used ecological traits as predictors (boosted regression trees, Bayesian additive regression trees, and neutral phylogeographic). A final hybrid model used a combination approach with a network backbone but incorporating the phylogeny and bat trait data. This eighth model was added in June 2020 as part of a collaborative open science program involving the use of preprints and public feedback.

All eight approaches were used to generate predictions about potential bat hosts of betacoronaviruses. A subset of five were used to recommend potential non-bat mammal hosts of betacoronaviruses (*k*-nearest neighbor, linear filtering, scaled-phylogeny, trait-free plug and play, and neutral phylogeographic). All of these methods were network-based except for neutral phylogeographic (Trait-3), which only uses simple phylogenetic and geographic predictors that have a consistent meaning across mammals and is trained on the entire mammal–virus network dataset rather than the bat–virus network. We did not use the other trait-based models to predict non-bat hosts. First, the ecological and morphological traits in PanTHERIA share less biological meaning at this scale (e.g., forearm length for bats, ungulates, and whales all have substantially different anatomical context). Second, assigning pseudoabsences to the vast majority (~3500 or more) of mammal species would create a severely unbalanced training dataset that would likely lead to poor predictions, given that only 109 betacoronavirus hosts were known during the first iteration of this analysis (79 bats and 30 other mammals).

*Network-based model 1: k-nearest neighbors recommender*

We follow the methodology previously developed for the recommendation of species feeding interactions^14^. This method builds a recommender system internally based on the *k*-nearest neighbor (kNN) algorithm, under which candidate hosts are recommended for a virus from a pool constituted by the hosts of the *k* viruses with which it has the greatest overlap (host sharing). Overlap is measured using the Jaccard-Tanimoto similarity, which is the cardinality of the intersection of two sets divided by the cardinality of their union. To obtain the pairwise similarity between two viruses, this divides the number of shared hosts by the cumulative number of hosts. The *k*NN of a virus are the *k* other viruses with which it has the highest Tanimoto similarity.

Hosts are then recommended by counting how many times they appear in these *k* neighbors, a quantity that ranges from 1 to *k*. We can impose arbitrary cutoffs by limiting the recommendations to the hosts that occur in at least *k*, *k-1*, etc, viruses. Previous leave-one-out validation of this model revealed that it is particularly effective for viruses with a reduced number of hosts^14^, which is likely to be the case for emerging viruses. Furthermore, the performance of this model was not significantly improved by the addition of functional traits, making it acceptable to run on the association data only.

This model was run two times: first, by measuring the similarity of viruses, and recommending hosts; second, by measuring the similarity of hosts, and recommending viruses. In all cases, only results for betacoronaviruses are reported, though the nearest neighbours are identified based on the entire network. The optimal value of *k* was determined by performing Leave-One-Out cross validation, wherein each interaction was set to 0, and we attempted to recover it with values of *k* ranging from 1 to 11, with a step of 1. The highest proportion of interactions correctly recovered peaked for *k*=5.

The outcome of this model should be subject to caution, as leave-one-out validation revealed that the success rate (i.e., ability to recover one interaction that has been removed) remained lower than 50% even when using *k*=5, and dropped as low as 5% when using *k*=1 (the nearest-neighbor algorithm). This strongly suggests that the dataset of reported host-virus associations is extremely incomplete; therefore, the identification of the nearest neighbors can be biased by under-reported interactions, and this can result in noise in the prediction. This noise can be particularly important when the *k*NN technique operates on viruses, of which the bat dataset has only 15.

*Network-based model 2: Linear filter recommender*

Following Stock *et al.**^15^*, we used a previously developed linear filter to infer potential missing interactions. This recommender system assumes that networks tend to be self-similar and uses this information to generate a score for an un-observed interaction that is a linear combination of the whether the interaction was observed (relative weight of 1/4), relative degree of host and virus, and the observed connectance of the network (all with relative weights of 1). As we are concerned with ranking interactions as opposed to examining the absolute value of a given score, the penalization coefficient associated to the interaction being presumed absent could be omitted with no change in the ranking, but has been set to a low value instead. The scores returned by the linear filter are not directly related to the probability of the interaction existing in this context, but higher scores still indicate interactions that are more likely to exist. Indeed, known hosts of betacoronavirus typically scored higher.

We used the zero-one-out approach to assess the performance of this model on the entire dataset. In all cases, non-interactions ranked lower than observed interactions even when entirely removing the penalization coefficient from the linear filter parameters, which suggests that the network structure (degree and connectance) is capturing information that describes which species can interact.

*Network-based model 3: Plug and play*

For network problems, the “plug and play” model is a statistical approach that formulates Bayes’ theorem for link prediction around the conditional density of traits of known associations compared to traits of every possible association in a network. That is, the probability of observing a host-virus association given the combination of host and viral traits for a given pair (P(y=1 | z), is equal to a combination of the trait space where an interaction occurs (f_1_(z)), the probability that a host-virus association occurs (P(Y=1)), and the available trait space of all possible host-virus associations (f(z)):

$$P(y=1|z)=(f_{1}(z)*P(Y=1))/f(z)$$

The conditional density function is measured by using non-parametric kernel density estimators (implemented with the R package *np*), and the conditional ratio between them is used to estimate link “suitability”, a scale-free ratio. Compared to other machine learning methods that fit training data iteratively, plug and play is comparatively simple, and directly infers the most likely extensions of observed patterns in data. The plug and play was originally developed to forecast missing links in host-parasite networks^16^, but has since been used to model species distributions^17^ and predict the global spread of human infectious diseases^18^. We used this model here to estimate suitability of host-virus interactions by first modeling the entire estimated network of host-virus interaction suitability, and ranking hosts that are not infected by betacoronaviruses by their estimated suitability for betacoronaviruses.

The “plug and play” model is trained using either matched pairs of host and pathogen ecological, morphological, or phylogenetic traits^16^, or by using a latent approach^18^ which considers the mean similarity of pathogens in their host ranges and the mean similarity of hosts in their pathogen communities as ‘traits’. We decided to use the latent approach, as host trait data was far more available than viral trait data. Further, the taxonomic scale considered for host (species) and virus (genus) differed, making the resolution of potential trait data different enough to potentially confound trait-based approaches in this modeling framework.

Relative suitability of a host-virus association, as estimated by the “plug and play” model, is formulated as a density ratio estimation problem. The suitability of a host-virus association is quantified as the quotient of the distribution of latent trait values when an association was recorded over the distribution of all the latent trait values. As an attempt to control for sampling effort of mammal and bat host species, we included PubMed citation counts for host species (as described above) in the estimation of host-virus suitability. We explored host-pathogen suitability using the entire mammal-virus associations dataset, to maximize the available information on the network’s structure, and ranked host-pathogen pairs by their relative suitability value. From the final predictions, we subset out bat-specific predictions. When predicting, we set citation counts to the mean of training data, as a sampling bias correction.

*Network-based model 4: Scaled-phylogeny*

We apply the network-based conditional model of Elmasri *et al.**^19^* for predicting missing links in bipartite ecological networks. The full model combines a hierarchical Bayesian latent score framework which accounts for the number of interactions per taxon, and a dependency among hosts based on evolutionary distances. To predict links based on evolutionary distance, the probability of a host-parasite interaction is taken as the sum of evolutionary distances to the documented hosts of that parasite. This allocates higher probabilities when a few closely related hosts, or many distantly related hosts interact with a parasite. In this way phylogenetic distances are combined with individual affinity parameters per taxa to model the conditional probability of an interaction.

In ecological studies, it is common to use time-scaled phylogenies to quantify evolutionary distance among species^20^. We may use these fixed evolutionary distances for link prediction, but parasite taxa are known to be more or less constrained by phylogenetic distances among hosts^21^. Further, phylogenies are hypotheses about evolutionary relationships and have uncertainties in the topology and relative distances among species^22^. Rather than treating phylogenetic distances as fixed, Elmasri *et al.**^19^* re-scale the phylogeny by applying a macroevolutionary model of trait evolution. While any evolutionary model that re-scales the phylogenetic distances among species may be used, we use the early-burst model, which allows evolutionary change to accelerate or decelerate through time^23^. This different emphasis is placed on deep versus recent host divergences when predicting links.

We apply the model to a network of associations among host species and viral genera, and the mammal supertree, which allows us to leverage information from across the network to predict undocumented bat-betacoronavirus associations. We fit sets of models, applying both the full model, and the phylogeny-only model to both the bat-viral genera associations, and the mammal-viral genera associations. For each data-model combination we fit the model using ten-fold cross-validation holding out links for which there is a minimum of two observed interactions. The posterior interaction matrices resulting from each of the ten models are then averaged to generate predictions for all links in the network, with betacoronaviruses subset to generate the ensemble predictions.

To assess predictive performance, we attempted to predict the withheld interactions, and calculated AUC scores by thresholding predicted probabilities per fold, and taking an average across the 10 folds. In addition to AUC, we also assessed the model based on the percent of documented interactions accurately recovered. For the bat-viral genera data the full model resulted in an average AUC of 0.82 and recovered an average of 90.1% of held out interactions, while the phylogeny-only model showed increased AUC (0.86), but a decreased proportion of held-out interactions recovered (84.5%). Interestingly, the models for bat-virus genera associations had marginally worse predictive performance compared to the same models run on the larger network of mammal-virus associations (full model: AUC 0.88, 84.4% positive interactions recovered; phylogeny-only model: AUC: 0.88, 88.8% positive interactions recovered), indicating that predicting bat-betacoronavirus associations may benefit from including data on non-bat hosts. The models also estimated the scaling parameter (eta) of the early-burst model to be positive (average eta=7.92 for the full model run on the bat subset), indicating accelerating evolution compared to the input tree (Supplemental Figure 5). This means that recent divergences are given more weight than deeper ones for determining bat-viral genera associations, which is consistent with recent work on viral sharing^5,24^.

*Trait-based model 1: Boosted regression trees*

Previous work has been highly successful in predicting zoonotic reservoirs using a combination of taxonomic, ecological, and geographic traits as predictors. This approach has been previously used to identify wildlife hosts of filoviruses^12,25^, flaviviruses^26,27^, henipaviruses^28^, *Borrelia burgdorferi**^29^*, to predict mosquito vectors of flaviviruses^30^, and to predict rodent reservoirs and tick vectors of zoonotic viruses^31,32^. These approaches treat the presence of a specific virus (or genus of viruses) or a zoonotic pathogen as an outcome variable, with negative values given for species not known to be hosts (pseudoabsences), and use machine learning to identify the characteristics that predispose animals to hosting pathogens of concern. By predicting the probability that a given pseudoabsence is a false negative, the method can infer potential undetected or undiscovered host species.

This approach has almost exclusively been implemented using boosted regression trees (BRT), a classification and regression tree (CART) machine learning method that became popular a decade ago for species distribution modeling.^33^ Boosted regression trees develop an ensemble of classification trees which iteratively explain the residuals of previous trees, up to a fixed tree depth (usually between 3 and 5 splits). The incorporation of boosting allows the model, as it is fit, to progressively better explain poorly-fit cases within training data.

We used boosted regression trees to identify trait profiles that predict bat hosts of betacoronaviruses, including all trait predictors from the trait database that met baseline coverage (< 50% missing values) and variation (< 97% homogenous) thresholds. For all model fitting, we specified a Bernoulli error distribution for our binary response variable and applied 10-fold cross validation to prevent overfitting (R package *gbm*). We started by fitting a global model to our full dataset, first specifying learning rate = 0.01 (shrinks the contribution of each tree to the model) and tree complexity = 4 (controls tree depth) as per default values and subsequently tuning to minimize cross-validation error.

We reduced the variable set by calling the gbm.simplify() function from the *dismo* package, which computes and compares the mean change in cross-validation error (deviance) produced by dropping different sets of least-contributing predictors^33^. The final simplified model included 23 variables, plus citation counts, which we added to correct for sampling bias.

We applied bootstrapping resampling methods to estimate uncertainty, using our tuned model to fit 1000 replicate models. For each model, training sets were assembled by randomly selecting with replacement 79 bat-coronavirus associations from the set of reported bat hosts and 79 pseudoabsences. Trained models were used to generate relative influence coefficients for trait predictors and coronavirus host probabilities across all bat species. Partial dependence plots display relative influence coefficients and bootstrapped confidence intervals for the top ten contributing trait predictors. The medians of host probabilities were ranked and used to identify the top ten candidate host species. When predicting, we set citation counts to the mean of training data, as a sampling bias correction.

*Trait-based model 2: Bayesian additive regression trees*

A similar workflow to trait-based model 1 was implemented using Bayesian additive regression trees (BART), an emerging machine learning tool that has similarities to more popular methods like random forests and boosted regression trees. BART adds several layers of methodological innovation, and performs well in comparison exercises with other advanced machine learning methods^34^. Several features make BART very convenient for modeling projects like these, including several easy-to-use implementations in R packages, built-in capacity to impute and predict on missing data, and easy construction of variable importance and partial dependence plots.

Like other classification and regression tree methods, BART assigns the probability of a binary outcome variable by developing a set of classification trees - in this case, a sum-of-trees model - that split data (“branches”) and assign values to terminal nodes (“leaves”). Whereas other similar methods generate uncertainty by adjusting data (e.g. random forests bootstrap training data and fit a tree to each bootstrap; boosted regression trees are usually implemented with iterated training-test splits to generate confidence intervals), BART characterizes uncertainty using an MCMC process. An initial sum-of-trees model is fit to the entire dataset, and then rulesets are adjusted in a limited and stochastic set of ways (e.g., adding a split; switching two internal nodes), with the sum-of-trees model backfit to each change. After a burn-in period, the cumulative set of sum-of-trees models is treated as a posterior distribution. This has some advantages over other methods, like boosted regression trees or random forests. In particular, posterior width is a direct response to model uncertainty (rather than approximating it by permuting training data), and a single model can be run (instead of an ensemble trained on smaller subsets of training data), allowing the model to use the full training dataset all at once.^35^

Unlike many Bayesian machine learning methods, BART is easily implemented out-of-the-box, due to a limited set of customization needs. Three main priors control the fitting process: one usually-uniform prior on variable importance, one two-parameter negative power distribution on tree depth (preventing overfitting), and an inverse chi-squared distribution on residual variance. A set of well-performing priors from the original BART study^34^ are widely used across R implementations for out-of-the-box settings, but can be further adjusted relative to modeling needs. In this study, we implemented BART models using a Dirichlet prior for variable importance (DART), a specification that is designed for situations with high dimensionality data that probably reflects a small number of true informative predictors. This often produces a much more reduced model without going through a stepwise variable selection process, which can be slow and very prone to stochasticity as well as highly subject to bias.^35,36^

We implemented this approach using the *BART* package in R, using the bat-virus association dataset to generate an outcome variable, and the bat traits dataset as predictors. BART models were implemented with 200 trees and 10,000 posterior draws, using every trait feature that was at least 50% complete and < 97% homogenous (taken from TBM1). The *BART* package implements an MCMC sampler that is readily used out-of-the-box, with a default of 100 burn-in samples and no other major aspects that require user-end input. Convergence diagnostics can be generated using elements of the model output (e.g., trace plots of the error variance). However, these diagnostics are largely unnecessary for the low-dimensional dataset used here, given other more informative ways of measuring model performance we implemented later.

We tried four total implementations, based on two decisions: BART uncorrected and corrected for citation counts (BART-u, BART-c), and DART uncorrected and corrected for citation counts (DART-u, DART-c). All four models performed well, with little variation in predictive power measured by the area under the receiver operator curve calculated on training data (BART-u: AUC = 0.93; BART-c: AUC = 0.93; DART-u: AUC = 0.93; DART-c: 0.90; Supplemental Figure 6). Across all models, spatial variables had a high importance, including some regionalization (extent of range) and some variables capturing larger geographic range sizes, as did a diet of invertebrates (pulling out the phylogenetic signal of insectivorous bats; Supplemental Figure 7).

All models identified a number of “false negative” hosts that would be suitable based on a 10% false negative classification threshold for known betacoronavirus hosts (implemented with the R package ‘PresenceAbsence’). BART-u identified 217 missing hosts, BART-c identified 279 missing hosts, DART-u identified 222 missing hosts, and DART-c identified 384 missing hosts, suggesting that this model most penalized overfitting as intended. As a result, we considered this model the most rigorous and powerful for inference and used DART-c in the final model ensemble. We predicted across all 1,040 bats without recorded betacoronavirus associations and ranked predicted probability. When predicting, we set citation counts to the mean of training data, as a sampling bias correction.

*Trait-based model 3: Phylogeographic neutral model*

We used a previously published pairwise viral sharing model^5^ to predict potential betacoronavirus hosts based on the sharing patterns of known hosts in a published dataset ^1^. We used a generalised additive mixed model (GAMM), which was fitted in the first half of 2019 using the *mgcv* package, with pairwise binary viral sharing (0/1 denoting if a species shares at least one virus) as a response variable. Explanatory variables include pairwise proportional phylogenetic distance and geographic range overlap (taken from the IUCN species ranges), with a multi-membership random effect to control for species-level sampling biases. The model was then used to predict the probability that a given species pair share at least one virus across 4,196 placental mammals with available data, producing a predicted viral sharing network that recapitulates a number of known macroecological patterns, as well as predicting reservoir hosts with surprising accuracy^5^. Subsetting this predicted sharing matrix, we listed the rank order of hosts most likely to share with all known betacoronavirus hosts in our datasets.

*Hybrid model 1: Two-step kernel ridge regression*

To make predictions using both the traits and the phylogeny of the hosts, we used Two-Step Kernel Ridge Regression (TSKRR), a kernel-based pairwise learning method^37–39^. This method can predict interactions between two species, i.e., hosts and viruses, both in-sample and out-of-sample. Conceptually, TSKRR performs nonlinear regression twice: once to generalize to new hosts and once to generalize to new viruses (Supplemental Figure 8A). As a kernel method, it can exploit arbitrary similarity measures to describe the species. At the same time, its conceptual simplicity permits using efficient shortcuts for tuning and specialized cross-validation.

For the hosts, we again restricted ourselves to bats. First, we imputed missing trait values using the *MissForest* package^40^. We then selected the relevant traits, standardized them, and computed the squared Euclidean distance. We plugged this distance into a standard radial basis kernel^41^, where we used the median heuristic to set the bandwidth. Similarly, for the bat phylogeny, we computed the distance matrix. We combined this distance with the radial basis kernel, again setting the bandwidth by the median of the distances. Bats were described using a kernel matrix derived from traits, phylogeny, or the average of both (Supplemental Figure 8B). We corrected for sampling bias by including the number of citations per species as a covariate and predicted using the average number of citations for all hosts. For viruses, we used a simple kernel with a “1” if two virus genera belong to the same family, and a “0” elsewise. For all the kernel matrices, we added 0.1 to each element and an additional 0.1 to the diagonal elements. This modification serves as an intercept and it gives the TSKRR the same capacity as the linear filtering method described earlier^37^.

We fit models using the R package *xnet* (<https://github.com/CenterForStatistics-UGent/xnet>). We assessed model performance in two ways. First, we used leave-one-out for every element of the incidence matrix and computed the AUC over all predicted interactions. Secondly, we used leave-one-out cross-validation on bats (i.e., leaving out a single species, making predictions for all the viruses of that bat, and repeating this for every species). In this setting, we computed the AUC for every virus over all bats and averaged these values (Supplemental Figure 8C). For both evaluation strategies, we removed viruses with no hosts in the dataset, although we kept these for training the model (Supplementary Table 4). Compared to the average performance for the viruses, betacoronaviruses were slightly more difficult to predict. Phylogeny was slightly more informative than traits, although the effect between them was synergistic. As would be expected, it was easier to predict missing interactions than to predict new hosts.

To assess which variables are most important, we computed feature importance based on random permutations. We randomly reshuffled one or both of the bats’ kernel matrices, destroying its information content. Subsequently, we recorded the decrease in AUC for the setting ‘interactions’ in the model as an indication of relative importance. We also randomly reshuffled each trait separately and monitored the effect on performance. We repeated each random permutation 100 times. Reshuffling the complete kernel matrix with both traits and phylogeny resulted in the largest average drop in AUC of 0.19 (Supplementary Figure 9). Reshuffling only the phylogeny kernel matrix had a much more profound impact than the trait matrix (0.08 vs. 0.05). Contributions of traits were smaller by comparison. Citation count was the largest effect (0.02), with diet breadth, population group size , and annual birth pulse as the remaining largest contributions (all <0.01). Removing citations resulted in a stronger importance for these variables but did not change the relative importance.

**Consensus Methods**

*Combining and ranking predictions*

For all eight models predicting bat hosts of betacoronaviruses, and five models predicting mammal hosts of betacoronaviruses, we combined predictions—generated using the same standardized data—into one standardized dataset. All mammal models were trained on data including bats, but predictions were subset to exclude bats to focus on likely intermediate hosts.

Each study’s unique output—a non-intercomparable mix of different definitions of suitability or probability of association—were transformed into proportional rank, where lower rank indicates higher evidence for association out of the total number of hosts examined. By rescaling all results to proportional ranks between zero and one, we also allowed comparison of in-sample and out-of-sample predictions across all models. Proportional ranks were averaged across models to generate one standardized list of predictions. This absorbed much of the variation in model performance (Supplemental Figure 1) and produced a set of rankings that performed well. This ensembling approach was selected as the simplest way of combining different models with different output formats. This was intended not to improve the overall performance higher than any given model but rather to develop the simplest possible set of consensus recommendations for use in field and laboratory sampling efforts, given initial uncertainty about the relative performance of any given model.

We elected not to withhold any “test” data to measure model performance, given that each method deployed in the ensemble has been independently and rigorously tested and validated in previous publications. Instead, to maximize the amount of available training data for every model, we used full datasets in each model and measured performance on the full training data.

For bats, our initial ensemble of models spanned a large range of performance on the training data, measured by the area under the receiver operator curve (AUC; Network 1: 0.624; Network 2: 0.987; Network 3: 0.514; Network 4: 0.726; Trait 1: 0.850; Trait 2: 0.902; Trait 3: 0.762; Hybrid 1: 0.924), indicating that it was possible to suitably detect differences in model performance on the full data. The total ensemble of proportional ranks performed adequately (AUC = 0.832). We used known betacoronavirus associations to threshold each model and the ensemble predictions based on a 10% omission threshold (90% sensitivity), and we again found a wide range in the number of predicted undiscovered bat hosts of betacoronaviruses (Network 1: 162 species; Network 2: 1; Network 3: 111; Network 4: 44; Trait 1: 425; Trait 2: 384; Trait 3: 720; Hybrid 1: 181; total ensemble: 239 species). Given concerns about mammal model performance and biological accuracy (see Main Text), we elected not to apply this exercise to mammal hosts at large.

*Geographic and taxonomic patterns from ensemble models*

To visualize the spatial distribution of predicted bat hosts from this initial ensemble, we used the IUCN Red List database of species geographic distributions. We took the 48 in-sample predictions and 323 out-of-sample predictions and combined these range maps to visualize species richness of all predicted hosts (Figure 2).

We used phylogenetic factorization to next flexibly identify taxonomic patterns in the consensus proportional rankings of likely bat hosts of betacoronaviruses. Phylogenetic factorization is a graph-partitioning algorithm that iteratively partitions a phylogeny in a series of generalized linear models to identify clades at any taxonomic level (e.g., rather than *a priori* comparing strictly among genera or family) that differ in a trait of interest^42^. Using the mammal supertree, we used the *phylofactor* package to partition proportional rank as a Gaussian-distributed variable. We determined the number of significant phylogenetic factors using a Holm's sequentially rejective 5% cutoff for the family-wise error rate. We applied this algorithm across our initial ensemble prediction datasets: in- and out-of-sample bat ranks.

To assess potential discrepancy between taxonomic patterns in model ensemble predictions and those of simply host betacoronavirus status itself, we ran a secondary phylogenetic factorization treating host status as a Bernoulli-distributed variable, with the same procedure applied to determine the number of significant phylogenetic factors (Supplemental Table 1).

**Betacoronavirus Data Update**

*Literature searches*

To locate new (or previously undetected) studies that identified betacoronaviruses in novel bat hosts, we regularly searched Google Scholar with broad search queries, including “betacoronavirus bat detection”, “novel bat betacoronavirus”, and “bat betacoronavirus metagenomic”, as well as other searches with terms in different combinations including “testing”, “isolation”, and “PCR”. Active data collection was supplemented by additional Google Scholar alerts for newly published studies that mentioned the top 10 in-sample and out-of-sample predicted hosts plus the term “betacoronavirus.” To confirm that no studies had gone undetected, we also reran the GenBank query on June 28, 2021 and confirmed that no additional samples had been deposited that included hosts in the metadata not previously identified by our initial search.

Over more than a year since the initial preprint and model ensemble, this led to the identification of 30 new hosts: *Hipposideros pomona, Hypsugo pulveratus, Scotophilus kuhlii, Myotis pequinius,* and *Myotis horsfieldii**^43^*; *Pteropus lylei**^44^*; *Hipposideros larvatus, Scotophilus heathii,* and *Hipposideros lekaguli**^45^*; *Desmodus rotundus**^46^**; Macroglossus minimus**^47^**; Hipposideros gigas**^48^**; Plecotus auritus* and *Tadarida teniotis**^49^*; *Artibeus jamaicensis* and *Carollia sowelli**^50^*; *Myonycteris angolensis, Nycteris macrotis,* and *Nanonycteris veldkampii**^51^*; *Pipistrellus deserti**^52^*; *Pipistrellus coromandra**^53^*; *Megaerops kusnotei**^54^*; *Rhinolophus shameli**^55^*; *Rhinolophus acuminatus**^56^*; *Nycteris gambiensis* and *Pteronotus personatus**^57^**; Rhinolophus malayanus* and *Rhinolophus stheno**^58^*, the former of which was absent from the initial GenBank scrape because the RmYN02 sample was not recorded as a *Betacoronavirus* at the time; *Vespertilio murinus**^59^*; and and finally *Myotis punicus,* which was deposited in GenBank in 2017 (accession MN823619), likely in connection with a related study on picornaviruses^60^.

*Metagenomics and transcriptomics*

We examined published metagenomic and transcriptomic datasets for evidence of betacoronavirus in suspected bat hosts. 232 sequencing libraries from 12 suspected bat hosts were identified from published mammalian short read datasets (NCBI Short Read Archive query (“Mammalia”[Organism] NOT “Homo sapiens”[Organism] NOT “Mus musculus”[Organism]) AND (“type_rnaseq”[Filter] OR “metagenomic”[Filter] OR “metatranscriptomic”[Filter]) AND “platform illumina”[Properties])^61^. We then performed a preliminary examination of libraries for evidence of betacoronaviruses using the Serratus platform^62^. Libraries with at least one hit to at least one region of a coronavirus genome were selected for downstream analysis, while libraries from studies involving experimental infections and libraries with hits only matching to a Patent or an unverified sequence were excluded. The remaining 52 libraries representing 8 bat species were downloaded from the European Nucleotide Archive and examined for genetic evidence of betacoronaviruses. Libraries were first trimmed and quality filtered^63,64^, then contigs were assembled using SPAdes v3.10.1^65^. Assembled contigs were analyzed by diamond blastx v0.8.20.82^66^ against the Genbank non-redundant protein database^61^. No published libraries contained evidence of betacoronaviruses. However, we note that these libraries were primarily generated for purposes other than viral discovery, so may not target appropriate tissues or use library preparation methods best suited to detecting viruses.

*The USAID PREDICT dataset*

As a final source of novel data, we explored the data generated by the USAID Emerging Pandemic Threats PREDICT program, which was newly released in its entirety in 2021. While many of the findings of the program have already been published, especially those that include the discovery of betacoronaviruses in bats, a number of novel viruses are reported in these data that have yet to be described in other scientific literature. We downloaded the publicly available PCR testing files from the USAID Development Data Library on June 25, 2021 (<https://data.usaid.gov/Global-Health-Security-in-Development-GHSD-/PREDICT-PCR-Tests/ganm-umas>), and aligned these data with the taxonomic information made available by the PREDICT program’s SpillOver Risk Ranking Tool (spillover.global; ^67^) to identify the genus associated with all reported bat coronaviruses. This led to the identification of 10 new hosts of betacoronaviruses: *Acerodon celebensis*, *Chaerephon pumilus*, *Epomops buettikoferi*, *Glauconycteris variegata*, *Hipposideros cervinus*, *Hipposideros fuliginosus*, *Megaerops ecaudatus*, *Myonycteris torquata*, *Neoromicia* cf. *somalicus*, and *Pteropus* cf. *conspicillatus*. These last two were taken as the valid host names, while noting the uncertainty in the host identification.

**Revised Ensemble**

Using the three sources of novel data on betacoronavirus hosts, we were able to compile a list of 40 newly identified bat hosts that were not included in our original training dataset. This allowed us to rerun the entire ensemble, including all eight component models, with an expanded copy of the dataset that included these 40 new hosts. We were also able to develop a new metric of model performance by evaluating the prevalence in the training data and the sensitivity evaluated on these 40 new hosts for each model at every threshold between 0 and 1. We call this the training prevalence-test sensitivity curve, and we used the area under this curve (i.e, AUC-TPTSC) as a measure of model performance. Values close to 0.5 indicate the model performs roughly as well as expected under random predictions, whereas values closer to 1 suggest the model is able to efficiently identify undiscovered hosts. We evaluated this metric for each of the eight models and both the original and updated ensemble. Using the AUC-TPTSC scores, we developed a weighted version of the updated ensemble, where ranks were subject to a weighted average based on the AUC-TPTSC of each model minus that of the lowest-performing model, incidentally eliminating the worst-performing model (Network-2).

We lastly identified geographic and taxonomic patterns in predictions from our weighted revised ensemble. We again used the IUCN Red List database of species geographic distributions to visualize the species richness of all 423 predicted hosts (Figure 5A). We then used phylogenetic factorization to flexibly identify taxonomic patterns in the proportional rankings of the weighted revised ensemble. In contrast to our analyses of proportional ranks from the initial ensemble (in which we analyzed species in- and out-of-sample separately), we applied phylogenetic factorization across the entire bat phylogeny.

**Insights into SARS-CoV-2’s emergence**

At the time of our initial ensemble in 2020, we attempted a secondary ensemble using five of our eight models to predict the broader mammal–virus network with a focus on potential betacoronavirus bridge hosts. At the time, only 30 non-bat hosts of betacoronaviruses were available in our data. However, we found poor concordance in predictions (Supplemental Figure 3). We instead focused on the outputs of Trait-3, which predicts how species should share viruses in nature based on their evolutionary history and geography.

*Rhinolophus-specific implementation of trait-based model 3*

We repeated Trait-3 focusing on virus sharing patterns of *Rhinolophus affinis* and *R. malayanus* specifically. Notable likely bridge host predictions included the hog badger *Arctonyx collaris* (Carnivora: Mustelidae), which was examined for SARS-CoV antibodies in 2003 and is reported in wildlife markets^68,69^; a selection of civet cats (Carnivora: Viverridae) including *Viverra* species*;* the binturong (*Arctitis binturong*)*;* and the masked palm civet (*Paguma larvata*), the latter of which were implicated in the chain of emergence for SARS-CoV^70,71^; and pangolins (Pholidota: Manidae) including *Manis javanica* and *Manis pentadactyla,* which have been hypothesised to be part of the emergence chain for SARS-CoV-2^72,73^.

Alongside these high-ranked species-level predictions, we visually examined how predictions varied across all mammal orders and families using the whole dataset (Supplemental Figure 4). Pangolins (Pholidota), treeshrews (Scandentia), carnivores (Carnivora), hedgehogs (Erinaceomorpha), and even-toed ungulates (Artiodactyla) had high mean predicted probabilities. Investigating family-level sharing probabilities revealed that civets (Viverridae) and mustelids (Mustelidae) were responsible for the high Carnivora probabilities, and mouse deer (Tragulidae) and bovids (Bovidae) were mainly responsible for high probabilities in the Artiodactyla (Supplemental Figure 4).

**Tables**

**Supplemental Table 1. Results of phylogenetic factorization applied to bat betacoronavirus positivity and predicted rank probabilities from our initial ensemble.** The number of retained phylogenetic factors (following a 5% family-wise error rate applied to GLMs), taxa corresponding to those clades, number of species, and either percent positive (betacoronavirus positivity) or mean predicted rank probabilities for the clade compared to the paraphyletic remainder are shown stratified by analyses of betacoronavirus data and models applied in- and out-of-sample.

| **Sample** | **Factor** | **Taxa** | **Tips** | **Clade** | **Other** |
| --- | --- | --- | --- | --- | --- |
| raw data | 1 | Noctilionoidea, Vespertilionoidea | 658 | 4.7% | 12.2% |
| in | 1 | Noctilionoidea, Vespertilionoidea | 160 | 0.550 | 0.393 |
| out | 1 | Noctilionidae, Mormoopidae, Phyllostomidae | 117 | 0.765 | 0.489 |
| out | 2 | Rhinolophidae | 48 | 0.265 | 0.547 |
| out | 3 | Emballonuridae, Craseonycteridae, Rhinopomatidae, Megadermatidae, Nycteridae, Hipposideridae, Hipposideridae, Myzopodidae, Thyropteridae, Furipteridae, Natalidae, Molossidae, Vespertilionidae | 498 | 0.539 | 0.515 |
| out | 4 | *Mosia, Emballonura, Coleura, Cyttarops, Diclidurus, Centronycteris, Cormura, Saccopteryx, Balantiopteryx, Peropteryx* | 29 | 0.754 | 0.521 |
| out | 5 | *Eonycteris, Macroglossus, Syconycteris, Notopteris, Melonycteris, Harpyionycteris, Dobsonia, Styloctenium, Neopteryx, Pteralopex, Acerodon, Pteropus* | 82 | 0.458 | 0.538 |
| out | 6 | Thyropteridae, Furipteridae, Natalidae | 10 | 0.836 | 0.526 |
| out | 7 | *Otomops, Promops, Molossus, Eumops* | 20 | 0.836 | 0.526 |

**Supplemental Table 2. Results of phylogenetic factorization applied to predicted rank probabilities of bat betacoronavirus hosts from our weighted revised ensemble.** The number of retained phylogenetic factors (following a 5% family-wise error rate applied to GLMs), taxa corresponding to those clades, number of species per clade, and mean predicted rank probabilities for the clade compared to the paraphyletic remainder are shown.

| **Factor** | **Taxa** | **Tips** | **Clade** | **Other** |
| --- | --- | --- | --- | --- |
| 1 | Mystacinidae, Noctilionidae, Mormoopidae, Phyllostomidae | 161 | 0.738 | 0.478 |
| 2 | *Mosia, Emballonura, Coleura, Rhynchonycteris, Cyttarops, Diclidurus, Centronycteris, Cormura, Saccopteryx, Balantiopteryx, Peropteryx* | 31 | 0.794 | 0.509 |
| 3 | Molossidae | 98 | 0.611 | 0.508 |
| 4 | Thyropteridae, Furipteridae, Natalidae | 12 | 0.867 | 0.514 |
| 5 | *Rousettus, Megaloglossus, Eidolon, Myonycteris, Plerotes, Casinycteris, Scotonycteris, Nanonycteris, Hypsignathus, Epomops, Micropteropus, Epomophorus* | 34 | 0.267 | 0.526 |
| 6 | *Sphaerias, Alionycteris, Otopteropus, Haplonycteris, Latidens, Penthetor, Thoopterus, Aethalops, Balionycteris, Chironax, Dyacopterus, Ptenochirus, Megaerops, Cynopterus* | 26 | 0.241 | 0.525 |
| 7 | Rhinolophidae | 73 | 0.345 | 0.531 |
| 8 | *Chaerephon, Mops* | 32 | 0.471 | 0.519 |

**Supplemental Table 3. Taxonomic scale of model training data and predictive implementation.** Notes: (1) These models generated predictions of sharing with *Rhinolophus affinis* over all non-human mammals in the HP3 dataset, then subsetted to bats. (2) In these models, bat-betacoronavirus predictions are based on a subset of binary outcomes for known association with betacoronaviruses, without any other viruses included.

| **Model approach** | **Training data scale** | **Bat**  ***Betacoronavirus* predictions** | **Mammal-wide**  ***Betacoronavirus* predictions** |
| --- | --- | --- | --- |
| **Network-based 1**  k-Nearest neighbors | Bat–virus | 🗸 |  |
| **Network-based 1**  k-Nearest neighbors | Mammal–virus |  | 🗸 |
| **Network-based 2**  Linear filter | Bat–-virus | 🗸 |  |
| **Network-based 2**  Linear filter | Mammal-virus |  | 🗸 |
| **Network-based 3**  Plug and play | Mammal–virus^1^ | 🗸 | 🗸 |
| **Network-based 4**  Scaled-phylogeny | Bat-virus | 🗸 |  |
| **Network-based 4**  Scaled-phylogeny | Mammal–virus |  | 🗸 |
| **Trait-based 1**  Boosted regression trees | Bat–betacoronavirus^2^ | 🗸 |  |
| **Trait-based 2**  Bayesian additive regression trees | Bat–betacoronavirus^2^ | 🗸 |  |
| **Trait-based 3**  Neutral phylogeographic | Mammal–virus^1^ | 🗸 | 🗸 |
| **Hybrid 1** Two-step kernel ridge regression | Bat–betacoronavirus | 🗸 |  |

**Supplemental Table 4. Model performance for the hybrid model.** Area under receiver-operator curve (AUC-ROC) for predicting interactions, or predicting for betacoronaviruses or for hosts, using bat traits, phylogeny or a combination of both.

|  | **interactions** | **Betacoronavirus** | **hosts (average)** |
| --- | --- | --- | --- |
| **traits** | 0.8784 | 0.6742 | 0.7304 |
| **phylogeny** | 0.8848 | 0.73389 | 0.7623 |
| **both** | 0.8975 | 0.7510 | 0.7909 |

**Supplemental Figure 1. Bat betacoronavirus model performance from initial analyses in 2020.** Curves show observed betacoronavirus hosts against predicted proportional ranks from eight individual models and those incorporated into one multi-model ensemble. Black lines show a binomial GLM fit to the predicted ranks against the recorded presence or absence of known betacoronavirus associations for in-sample hosts. Points are jittered to reduce overlap.

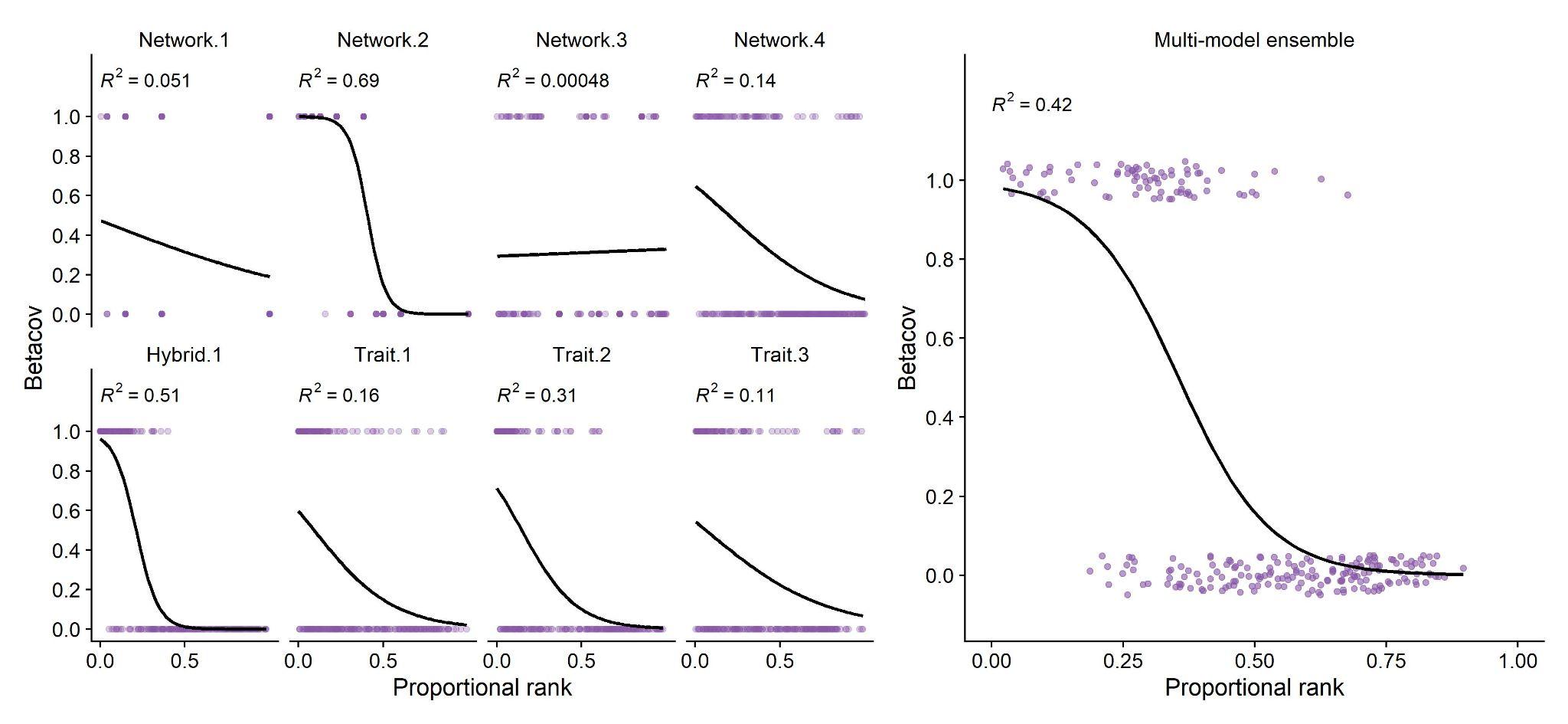

**Supplemental Figure 2. Bat betacoronavirus model performance from our revised 2021 ensemble after incorporating 40 new host species.** Curves show observed betacoronavirus hosts against predicted proportional ranks from eight individual models, and those incorporated into one multi-model ensemble. Black lines show a binomial GLM fit to the predicted ranks against the recorded presence or absence of known betacoronavirus associations for in-sample hosts. Points are jittered to reduce overlap.

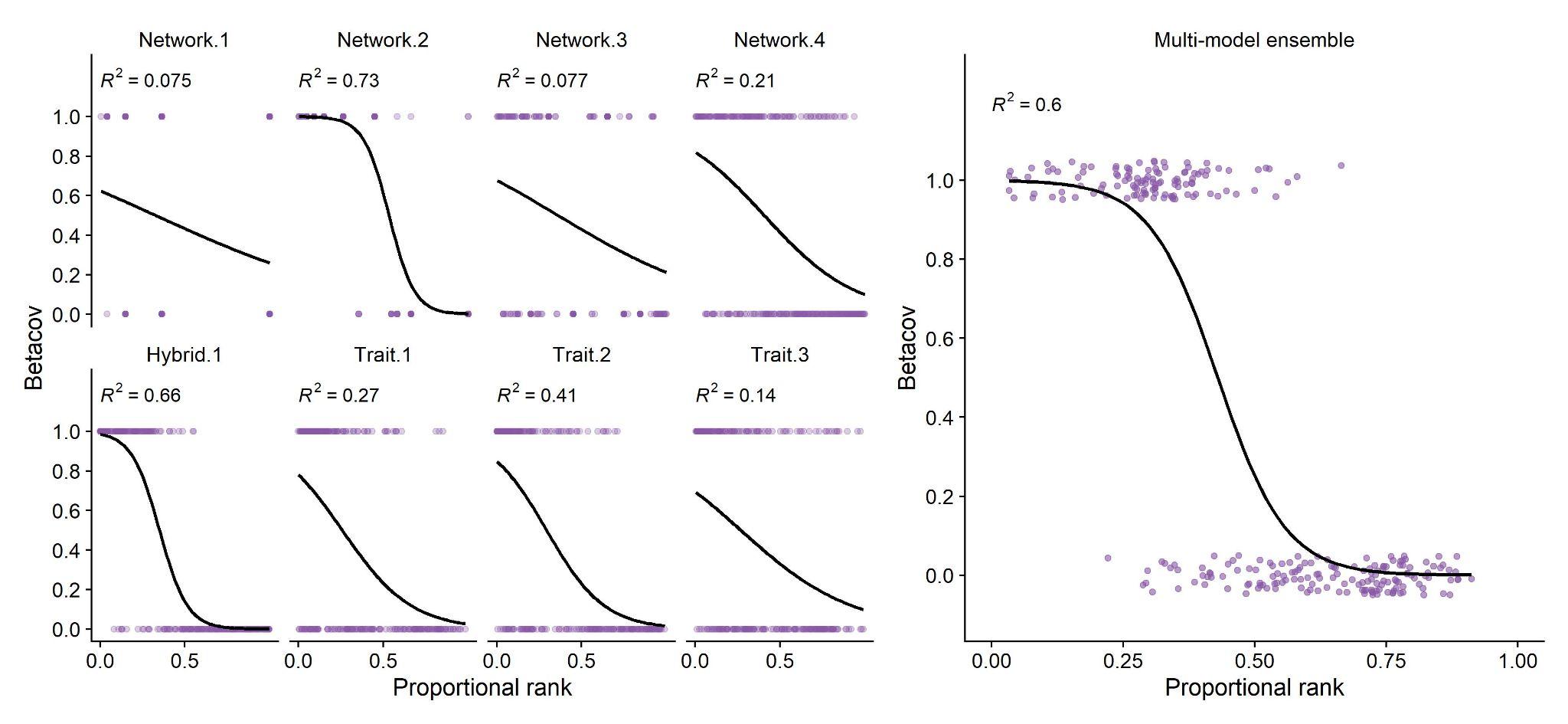

**Supplemental Figure 3. Poor concordance among predictive models for mammal hosts of betacoronaviruses.** The pairwise Spearman’s rank correlations between models’ ranked species-level predictions were generally low (A). In-sample predictions varied significantly and heavily prioritized domestic animals and well-studied hosts (B). The ten species with the highest mean proportional ranks across all models are highlighted in shades of purple. Only in-sample predictions are displayed because only one model (Trait-based 3) was able to predict out of sample for all mammals.

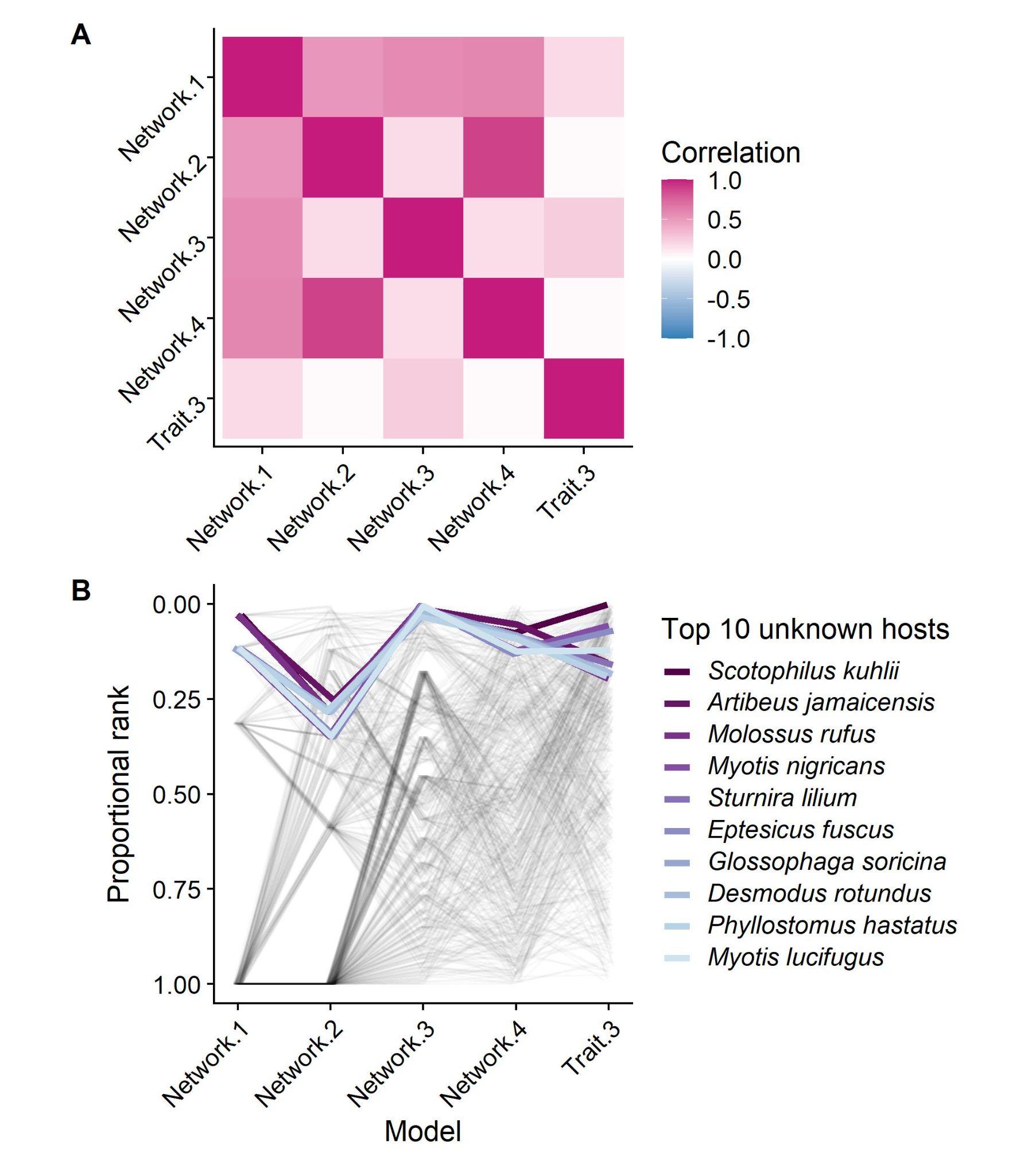

**Supplemental Figure 4.** Predicted species-level sharing probabilities of A) *Rhinolophus affinis* and B) *Rhinolophus malayanus*, calculated according to the phylogeographic viral sharing model^. Each coloured point is a mammal species. Black points and error bars denote means and standard errors for each order. Mammal families are arranged according to their mean sharing probability.

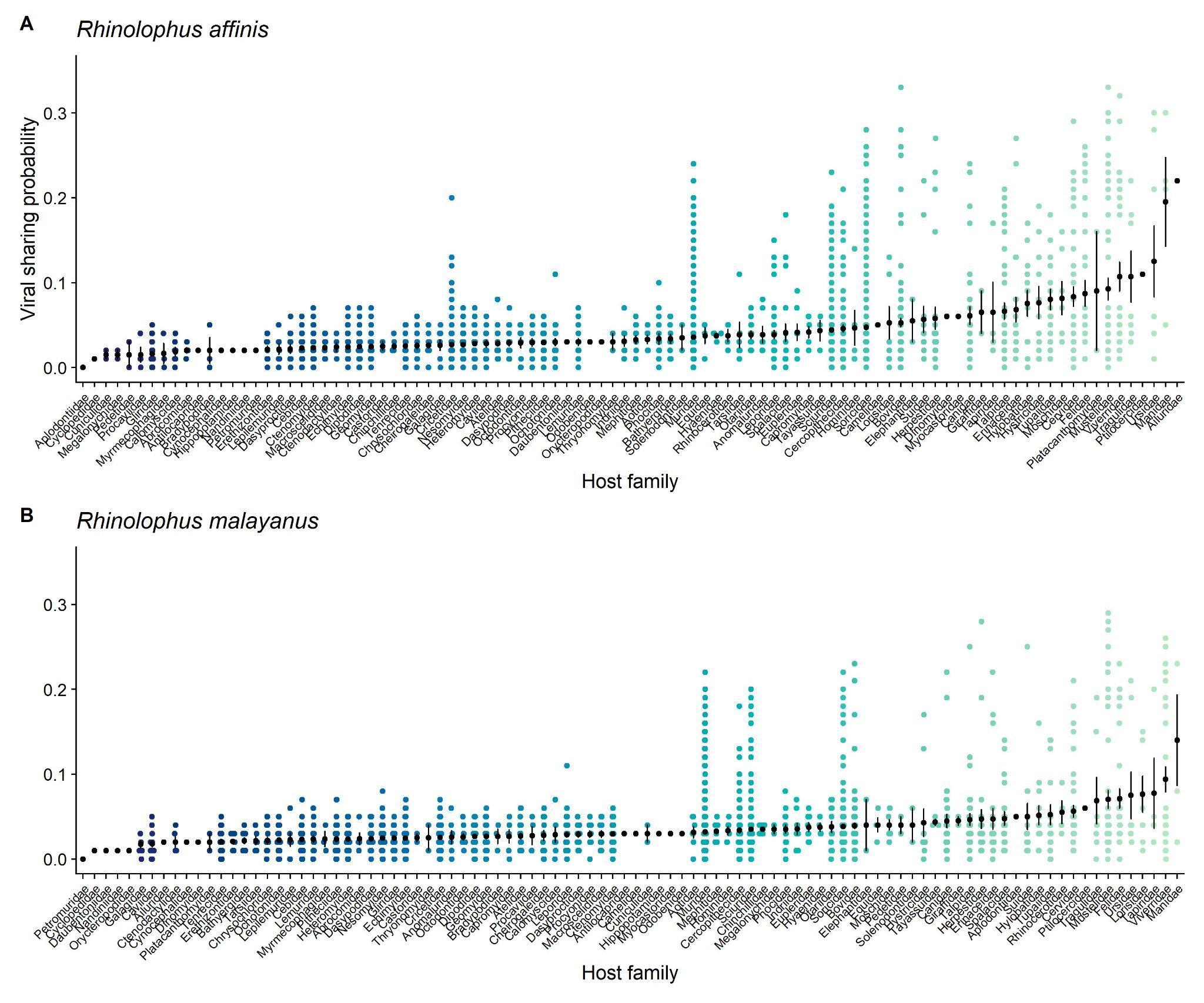

**Supplemental Figure 5.** To account for uncertainty in the phylogenetic distances among hosts, the scaled-phylogeny model estimates a tree scaling parameter (eta) based on an early-burst model of evolution. On the left is the unscaled bat phylogeny for the hosts in the bat-virus genera network, and on the right is the same tree rescaled according to mean estimated scaling parameter (eta = 7.92). Eta values above 1 indicate accelerating evolution, suggesting less phylogenetic conservatism in host-virus associations among closely related taxa than would be predicted by a Brownian-motion model on the unscaled tree.

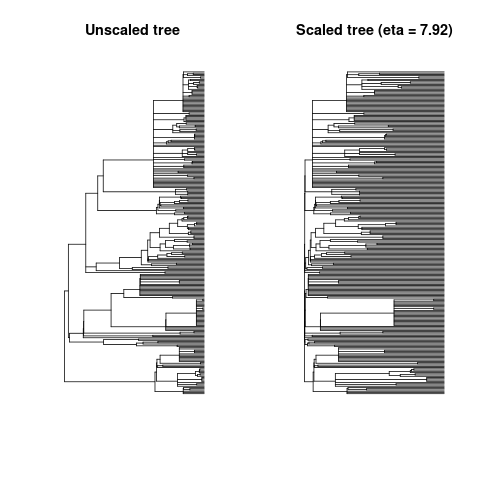

**Supplemental Figure 6.** Four formulations of Bayesian additive regression tree (BART) models produce slightly different results, but largely agree. Two models use baseline BART, while two models use a Dirichlet prior on variable importance (DART). Two are uncorrected for sampling bias (u) while two are corrected using citation counts (c). In the final main-text model ensemble, we present a DART model including correction for citation bias, which penalizes overfitting and spurious patterns two ways and leads to predictions with a lower total correlation with the data, but a still-high performance (AUC = 0.90).

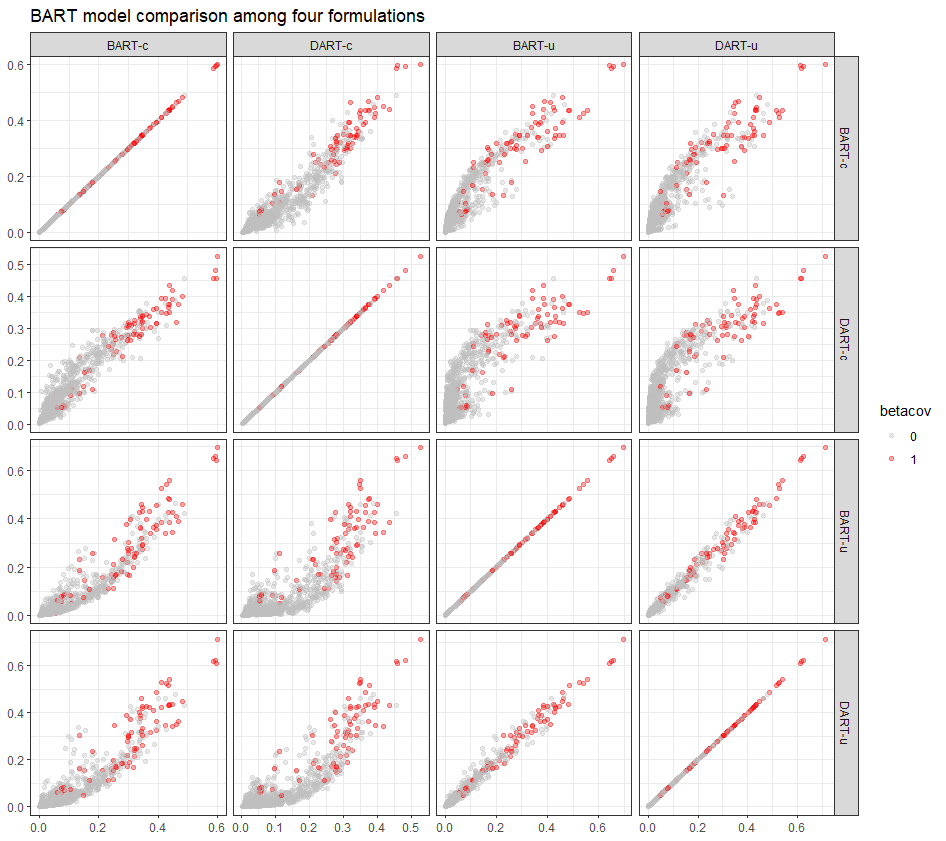

**Supplemental Figure 7.** Variable importance plots for the Bayesian additive regression tree models with uniform variable importance prior (top) versus Dirichlet prior (bottom), without (left) and with (right) correction for citations.

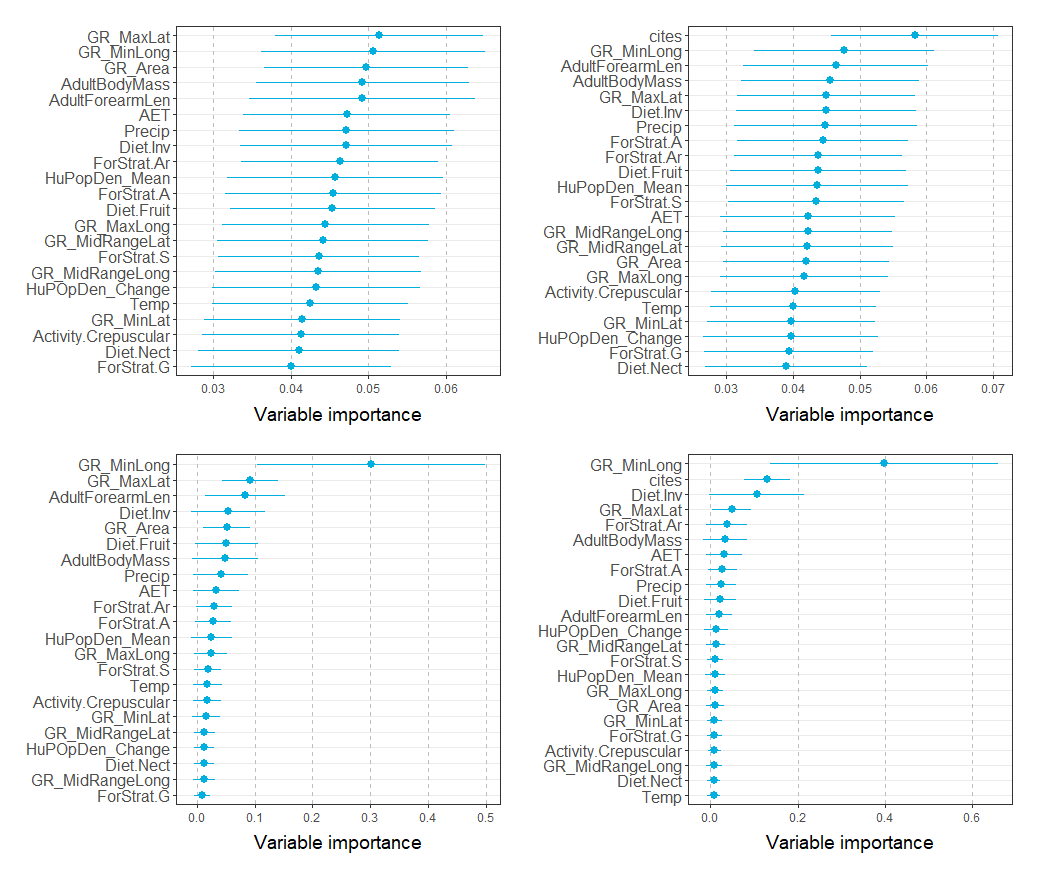

**Supplemental Figure 8. Overview of the Two-Step Kernel Ridge Regression method.** A. The incidence matrix *Y* is modelled by two kernel matrices *K* and *G*, describing the hosts (here: bats) and viruses, respectively. The equations for fitting the model and making predictions are given below. B. Computing the host kernel matrix is done based on traits (*T)*, phylogeny (*P*), or both. A standard radial basis kernel is used to transform a distance (*d_ij_)* to a valid kernel. The two kernels are combined by averaging. C. The models are validated by specialized cross-validation. The setting interactions leaves one interaction out at a time and repeats this for every interaction. AUC is computed in micro-fashion. In the setting hosts, we leave out one host at a time and compute the AUC per virus, averaging the individual results. D. To assess the variable importance, we first randomly reshuffle the kernel matrices and monitor how the performance of a model fitted on these data drops. In a more fine-grained approach, we just reshuffle a single trait (indicated by the red arrow) and compare the resulting performances.

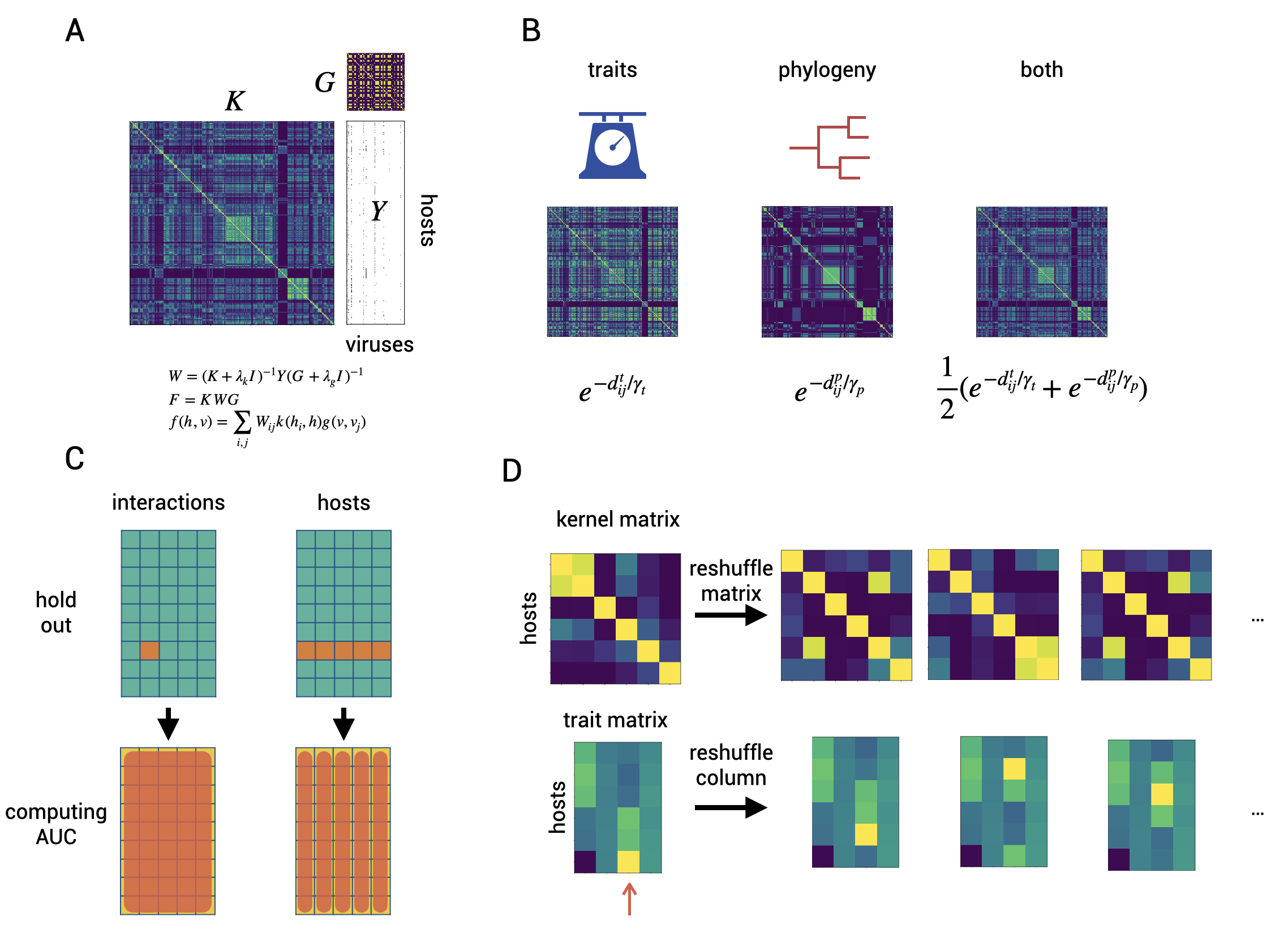

**Supplemental Figure 9. Illustration of the TSKRR model and the variable importance scores.** A**.** Heatmap of the prediction scores, together with a hierarchical clustering based on the kernel distance of *K* and *G*. B. Variable importance computed by random permutations of the kernel matrices or a single trait.

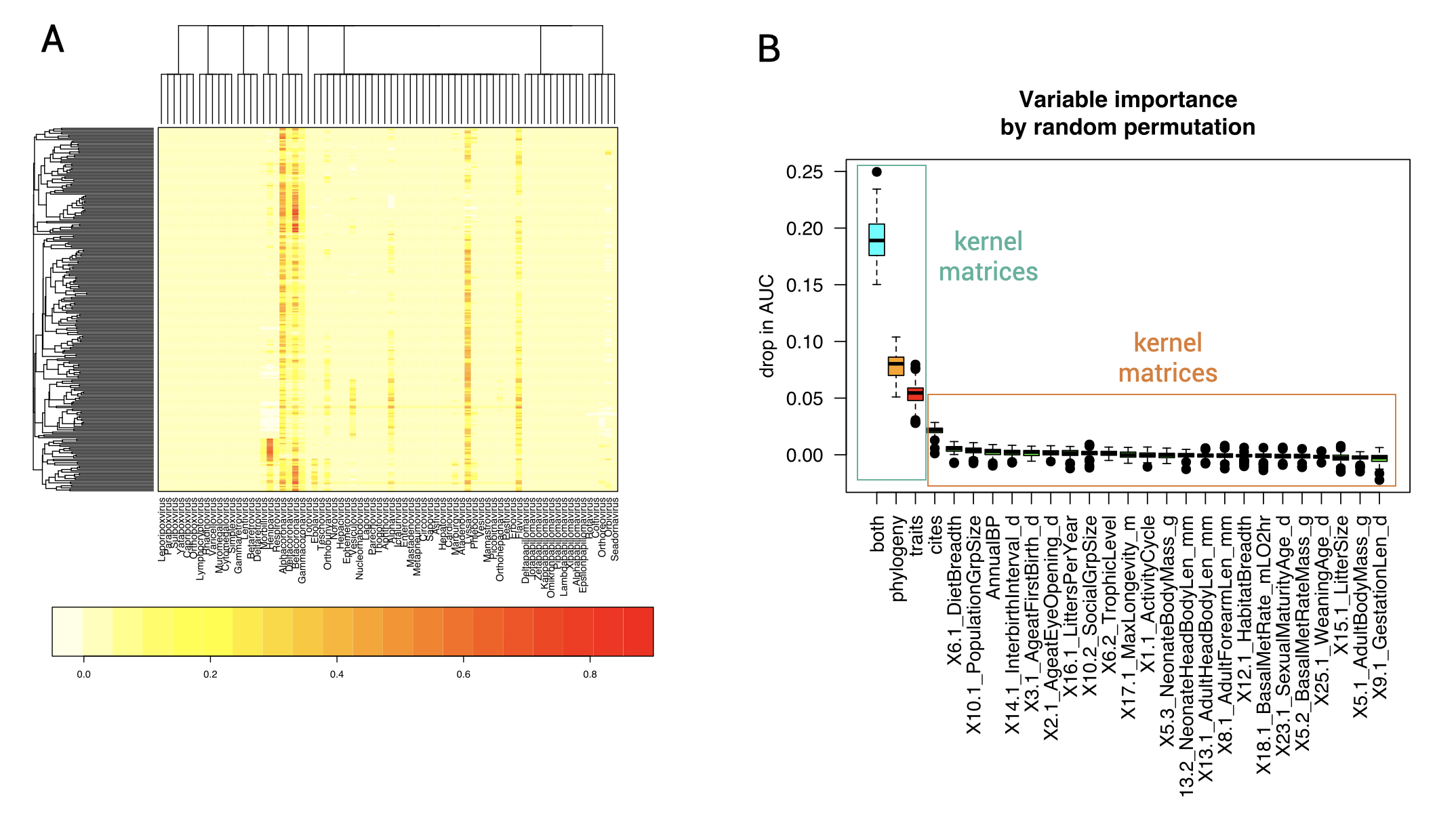

**Bibliography**

1	Olival KJ, Hosseini PR, Zambrana-Torrelio C, Ross N, Bogich TL, Daszak P. Host and viral traits predict zoonotic spillover from mammals. *Nature* 2017; **546**: 646–50.

2	Gibb R, Albery GF, Becker DJ, *et al.* Data proliferation, reconciliation, and synthesis in viral ecology. Cold Spring Harbor Laboratory. 2021; : 2021.01.14.426572.

3	Guth S, Visher E, Boots M, Brook CE. Host phylogenetic distance drives trends in virus virulence and transmissibility across the animal–human interface. *Philos Trans R Soc Lond B Biol Sci* 2019; **374**: 20190296.

4	Washburne AD, Crowley DE, Becker DJ, *et al.* Taxonomic patterns in the zoonotic potential of mammalian viruses. *PeerJ* 2018; **6**: e5979.

5	Albery GF, Eskew EA, Ross N, Olival KJ. Predicting the global mammalian viral sharing network using phylogeography. *Nat Commun* 2020; **11**: 2260.

6	Fritz SA, Bininda-Emonds ORP, Purvis A. Geographical variation in predictors of mammalian extinction risk: big is bad, but only in the tropics. *Ecol Lett* 2009; **12**: 538–49.

7	Redondo RAF, Brina LPS, Silva RF, Ditchfield AD, Santos FR. Molecular systematics of the genus Artibeus (Chiroptera: Phyllostomidae). *Mol Phylogenet Evol* 2008; **49**: 44–58.

8	Bouchard S. Chaerephon pumilus. *Mammalian Species* 1998; : 1–6.

9	Hoofer SR, Van Den Bussche RA, HoráČek I. Generic Status of the American Pipistrelles (Vespertilionidae) with Description of a New Genus. *J Mammal* 2006; **87**: 981–92.

10	Jones KE, Bielby J, Cardillo M, *et al.* PanTHERIA: a species-level database of life history, ecology, and geography of extant and recently extinct mammals. Ecology. 2009; **90**: 2648–2648.

11	Wilman H, Belmaker J, Simpson J, de la Rosa C, Rivadeneira MM, Jetz W. EltonTraits 1.0: Species-level foraging attributes of the world’s birds and mammals. Ecology. 2014; **95**: 2027–2027.

12	Han BA, Schmidt JP, Alexander LW, Bowden SE, Hayman DTS, Drake JM. Undiscovered Bat Hosts of Filoviruses. *PLoS Negl Trop Dis* 2016; **10**: e0004815.

13	Norberg A, Abrego N, Guillaume Blanchet F, *et al.* A comprehensive evaluation of predictive performance of 33 species distribution models at species and community levels. Ecological Monographs. 2019; **89**. DOI:[10.1002/ecm.1370](http://dx.doi.org/10.1002/ecm.1370).

14	Desjardins-Proulx P, Laigle I, Poisot T, Gravel D. Ecological interactions and the Netflix problem. *PeerJ* 2017; **5**: e3644.

15	Stock M, Poisot T, Waegeman W, De Baets B. Linear filtering reveals false negatives in species interaction data. *Sci Rep* 2017; **7**: 45908.

16	Dallas T, Park AW, Drake JM. Predicting cryptic links in host-parasite networks. *PLoS Comput Biol* 2017; **13**: e1005557.

17	Drake JM, Richards RL. Estimating environmental suitability. Ecosphere. 2018; **9**: e02373.

18	Dallas TA, Carlson CJ, Poisot T. Testing predictability of disease outbreaks with a simple model of pathogen biogeography. *R Soc Open Sci* 2019; **6**: 190883.

19	Elmasri M, Farrell MJ, Jonathan Davies T, Stephens DA. A hierarchical Bayesian model for predicting ecological interactions using scaled evolutionary relationships. The Annals of Applied Statistics. 2020; **14**: 221–40.

20	Cadotte MW, Jonathan Davies T, Regetz J, Kembel SW, Cleland E, Oakley TH. Phylogenetic diversity metrics for ecological communities: integrating species richness, abundance and evolutionary history. *Ecol Lett* 2010; **13**: 96–105.

21	Park AW, Farrell MJ, Schmidt JP, *et al.* Characterizing the phylogenetic specialism–generalism spectrum of mammal parasites. Proceedings of the Royal Society B: Biological Sciences. 2018; **285**: 20172613.

22	Harvey PH, Pagel MD. The comparative method in evolutionary biology. Oxford University Press, USA, 1998.

23	Harmon LJ, Losos JB, Jonathan Davies T, *et al.* Early bursts of body size and shape evolution are rare in comparative data. *Evolution* 2010; **64**: 2385–96.

24	Mollentze N, Streicker DG. Viral zoonotic risk is homogenous among taxonomic orders of mammalian and avian reservoir hosts. *Proc Natl Acad Sci U S A* 2020; **117**: 9423–30.

25	Schmidt JP, Maher S, Drake JM, Huang T, Farrell MJ, Han BA. Ecological indicators of mammal exposure to Ebolavirus. *Philos Trans R Soc Lond B Biol Sci* 2019; **374**: 20180337.

26	Han BA, Majumdar S, Calmon FP, *et al.* Confronting data sparsity to identify potential sources of Zika virus spillover infection among primates. *Epidemics* 2019; **27**: 59–65.

27	Pandit PS, Doyle MM, Smart KM, Young CCW, Drape GW, Johnson CK. Predicting wildlife reservoirs and global vulnerability to zoonotic Flaviviruses. *Nat Commun* 2018; **9**: 5425.

28	Plowright RK, Becker DJ, Crowley DE, *et al.* Prioritizing surveillance of Nipah virus in India. *PLoS Negl Trop Dis* 2019; **13**: e0007393.

29	Becker DJ, Han BA. The macroecology and evolution of avian competence for Borrelia burgdorferi. DOI:[10.1101/2020.04.15.040352](http://dx.doi.org/10.1101/2020.04.15.040352).

30	Evans MV, Dallas TA, Han BA, Murdock CC, Drake JM. Data-driven identification of potential Zika virus vectors. *Elife* 2017; **6**. DOI:[10.7554/eLife.22053](http://dx.doi.org/10.7554/eLife.22053).

31	Han BA, Schmidt JP, Bowden SE, Drake JM. Rodent reservoirs of future zoonotic diseases. Proceedings of the National Academy of Sciences. 2015; **112**: 7039–44.

32	Yang LH, Han BA. Data-driven predictions and novel hypotheses about zoonotic tick vectors from the genus Ixodes. *BMC Ecol* 2018; **18**: 7.

33	Elith J, Leathwick JR, Hastie T. A working guide to boosted regression trees. *J Anim Ecol* 2008; **77**: 802–13.

34	Chipman HA, George EI, McCulloch RE. BART: Bayesian additive regression trees. The Annals of Applied Statistics. 2010; **4**: 266–98.

35	Carlson CJ. embarcadero: Species distribution modelling with Bayesian additive regression trees in R. DOI:[10.1101/774604](http://dx.doi.org/10.1101/774604).

36	Whittingham MJ, Stephens PA, Bradbury RB, Freckleton RP. Why do we still use stepwise modelling in ecology and behaviour? *J Anim Ecol* 2006; **75**: 1182–9.

37	Stock M, Pahikkala T, Airola A, De Baets B, Waegeman W. A Comparative Study of Pairwise Learning Methods Based on Kernel Ridge Regression. *Neural Comput* 2018; **30**: 2245–83.

38	Stock M, Pahikkala T, Airola A, Waegeman W, De Baets B. Algebraic shortcuts for leave-one-out cross-validation in supervised network inference. *Brief Bioinform* 2018; published online Oct 16. DOI:[10.1093/bib/bby095](http://dx.doi.org/10.1093/bib/bby095).

39	Stock M, Piot N, Vanbesien S, Meys J, Smagghe G, De Baets B. Pairwise learning for predicting pollination interactions based on traits and phylogeny. *Ecol Modell* 2021; **451**: 109508.

40	Stekhoven DJ, Bühlmann P. MissForest—non-parametric missing value imputation for mixed-type data. *Bioinformatics* 2012; **28**: 112–8.

41	Schölkopf B, Smola AJ, Managing Director of the Max Planck Institute for Biological Cybernetics in Tubingen Germany Profe Bernhard Scholkopf, Bach F. Learning with Kernels: Support Vector Machines, Regularization, Optimization, and Beyond. MIT Press, 2002.

42	Washburne AD, Silverman JD, Morton JT, *et al.* Phylofactorization: a graph partitioning algorithm to identify phylogenetic scales of ecological data. *Ecol Monogr* 2019; **89**: e01353.

43	Latinne A, Hu B, Olival KJ, *et al.* Origin and cross-species transmission of bat coronaviruses in China. DOI:[10.1101/2020.05.31.116061](http://dx.doi.org/10.1101/2020.05.31.116061).

44	Wacharapluesadee S, Duengkae P, Chaiyes A, *et al.* Longitudinal study of age-specific pattern of coronavirus infection in Lyle’s flying fox (Pteropus lylei) in Thailand. *Virol J* 2018; **15**: 38.

45	Wacharapluesadee S, Duengkae P, Rodpan A, *et al.* Diversity of coronavirus in bats from Eastern Thailand. *Virol J* 2015; **12**: 57.

46	Brandão PE, Scheffer K, Villarreal LY, *et al.* A coronavirus detected in the vampire bat Desmodus rotundus. *Braz J Infect Dis* 2008; **12**: 466–8.

47	Tampon NVT, Rabaya YMC, Malbog KMA. First Molecular Evidence for Bat Betacoronavirus in Mindanao. *Philippine Journal of* 2020. <https://www.researchgate.net/profile/Marion-John-Michael-Achondo/publication/338817325_First_Molecular_Evidence_for_Bat_Betacoronavirus_in_Mindanao/links/5e2bc49e299bf152167b38de/First-Molecular-Evidence-for-Bat-Betacoronavirus-in-Mindanao.pdf>.

48	Maganga GD, Pinto A, Mombo IM, *et al.* Genetic diversity and ecology of coronaviruses hosted by cave-dwelling bats in Gabon. *Sci Rep* 2020; **10**: 7314.

49	Lecis R, Mucedda M, Pidinchedda E, Pittau M, Alberti A. Molecular identification of Betacoronavirus in bats from Sardinia (Italy): first detection and phylogeny. Virus Genes. 2019; **55**: 60–7.

50	Colunga-Salas P, Hernández-Canchola G. Bats and humans during the SARS-CoV-2 outbreak: The case of bat-coronaviruses from Mexico. *Transbound Emerg Dis* 2021; **68**: 987–92.

51	Lacroix A, Vidal N, Keita AK, *et al.* Wide Diversity of Coronaviruses in Frugivorous and Insectivorous Bat Species: A Pilot Study in Guinea, West Africa. *Viruses* 2020; **12**. DOI:[10.3390/v12080855](http://dx.doi.org/10.3390/v12080855).

52	Shehata MM, Chu DKW, Gomaa MR, *et al.* Surveillance for Coronaviruses in Bats, Lebanon and Egypt, 2013–2015. Emerging Infectious Diseases. 2016; **22**: 148–50.

53	Lacroix A, Duong V, Hul V, *et al.* Genetic diversity of coronaviruses in bats in Lao PDR and Cambodia. *Infect Genet Evol* 2017; **48**: 10–8.

54	Xu L, Zhang F, Yang W, *et al.* Detection and characterization of diverse alpha- and betacoronaviruses from bats in China. *Virol Sin* 2016; **31**: 69–77.

55	Hul V, Delaune D, Karlsson EA, *et al.* A novel SARS-CoV-2 related coronavirus in bats from Cambodia. DOI:[10.1101/2021.01.26.428212](http://dx.doi.org/10.1101/2021.01.26.428212).

56	Wacharapluesadee S, Tan CW, Maneeorn P, *et al.* Evidence for SARS-CoV-2 related coronaviruses circulating in bats and pangolins in Southeast Asia. *Nat Commun* 2021; **12**: 972.

57	Olival KJ, Cryan PM, Amman BR, *et al.* Possibility for reverse zoonotic transmission of SARS-CoV-2 to free-ranging wildlife: A case study of bats. *PLoS Pathog* 2020; **16**: e1008758.

58	Zhou H, Ji J, Chen X, *et al.* Identification of novel bat coronaviruses sheds light on the evolutionary origins of SARS-CoV-2 and related viruses. *Cell* 2021; published online June 9. DOI:[10.1016/j.cell.2021.06.008](http://dx.doi.org/10.1016/j.cell.2021.06.008).

59	Hardmeier I, Aeberhard N, Qi W, *et al.* Metagenomic analysis of fecal and tissue samples from 18 endemic bat species in Switzerland revealed a diverse virus composition including potentially zoonotic viruses. *PLoS One* 2021; **16**: e0252534.

60	Zeghbib S, Herczeg R, Kemenesi G, *et al.* Genetic characterization of a novel picornavirus in Algerian bats: co-evolution analysis of bat-related picornaviruses. *Sci Rep* 2019; **9**: 15706.

61	NCBI Resource Coordinators. Database resources of the National Center for Biotechnology Information. *Nucleic Acids Res* 2018; **46**: D8–13.

62	Edgar RC, Taylor J, Lin V, *et al.* Petabase-scale sequence alignment catalyses viral discovery. DOI:[10.1101/2020.08.07.241729](http://dx.doi.org/10.1101/2020.08.07.241729).

63	Martin M. Cutadapt removes adapter sequences from high-throughput sequencing reads. EMBnet.journal. 2011; **17**: 10.

64	Schmieder R, Edwards R. Quality control and preprocessing of metagenomic datasets. Bioinformatics. 2011; **27**: 863–4.

65	Bankevich A, Nurk S, Antipov D, *et al.* SPAdes: A New Genome Assembly Algorithm and Its Applications to Single-Cell Sequencing. Journal of Computational Biology. 2012; **19**: 455–77.

66	Buchfink B, Xie C, Huson DH. Fast and sensitive protein alignment using DIAMOND. Nature Methods. 2015; **12**: 59–60.

67	Grange ZL, Goldstein T, Johnson CK, *et al.* Ranking the risk of animal-to-human spillover for newly discovered viruses. *Proceedings of the National Academy of Sciences* 2021; : in press.

68	Guan Y, Zheng BJ, He YQ, *et al.* Isolation and characterization of viruses related to the SARS coronavirus from animals in southern China. *Science* 2003; **302**: 276–8.

69	Chen W, Newman C, Liu Z, *et al.* The illegal exploitation of hog badgers (Arctonyx collaris) in China: genetic evidence exposes regional population impacts. Conservation Genetics Resources. 2015; **7**: 697–704.

70	Wang M, Yan M, Xu H, *et al.* SARS-CoV Infection in a Restaurant from Palm Civet. Emerging Infectious Diseases. 2005; **11**: 1860–5.

71	Song H-D, Tu C-C, Zhang G-W, *et al.* Cross-host evolution of severe acute respiratory syndrome coronavirus in palm civet and human. *Proc Natl Acad Sci U S A* 2005; **102**: 2430–5.

72	Lam TT-Y, Shum MH-H, Zhu H-C, *et al.* Identifying SARS-CoV-2 related coronaviruses in Malayan pangolins. *Nature* 2020; published online March 26. DOI:[10.1038/s41586-020-2169-0](http://dx.doi.org/10.1038/s41586-020-2169-0).

73	Xiao K, Zhai J, Feng Y, *et al.* Isolation of SARS-CoV-2-related coronavirus from Malayan pangolins. *Nature* 2020; published online May 7. DOI:[10.1038/s41586-020-2313-x](http://dx.doi.org/10.1038/s41586-020-2313-x).
